# COLLAPSE: A representation learning framework for identification and characterization of protein structural sites

**DOI:** 10.1101/2022.07.20.500713

**Authors:** Alexander Derry, Russ B. Altman

## Abstract

The identification and characterization of the structural sites which contribute to protein function are crucial for understanding biological mechanisms, evaluating disease risk, and developing targeted therapies. However, the quantity of known protein structures is rapidly outpacing our ability to functionally annotate them. Existing methods for function prediction either do not operate on local sites, suffer from high false positive or false negative rates, or require large site-specific training datasets, necessitating the development of new computational methods for annotating functional sites at scale. We present COLLAPSE (Compressed Latents Learned from Aligned Protein Structural Environments), a framework for learning deep representations of protein sites. COLLAPSE operates directly on the 3D positions of atoms surrounding a site and uses evolutionary relationships between homologous proteins as a self-supervision signal, enabling learned embeddings to implicitly capture structure-function relationships within each site. Our representations generalize across disparate tasks in a transfer learning context, achieving state-of-the-art performance on standardized benchmarks (protein-protein interactions and mutation stability) and on the prediction of functional sites from the Prosite database. We use COLLAPSE to search for similar sites across large protein datasets and to annotate proteins based on a database of known functional sites. These methods demonstrate that COLLAPSE is computationally efficient, tunable, and interpretable, providing a general-purpose platform for computational protein analysis.

## 1. Introduction

The three-dimensional structure of a protein determines its functional characteristics and ability to interact with other molecules, including other proteins, endogenous small molecules, and therapeutic drugs. Biochemical interactions occur at specific regions of the protein known as functional sites. We consider functional sites that range from a few atoms which coordinate an ion or catalyze a reaction to larger regions which binds a cofactor or form a protein-protein interaction surface. The identification of such sites—and accurate modeling of the local structure-function relationship—is critical for determining a protein’s biological role, including our understanding of disease pathogenesis and ability to develop targeted therapies or protein engineering technologies. Significant effort has gone into curating databases to catalog these structure-function relationships, ^1–3^ but this cannot keep up with the rapid increase in proteins in need of annotation. The number of proteins of the Protein Data Bank (PDB) ^4^ increases each year, and AlphaFold ^5^ has added high-quality predicted structures for hundreds of thousands more. This explosion of protein structure data necessitates the development of computational methods for identifying, characterizing, and comparing functional sites at proteome scale.

Many widely used methods for protein function identification are based on sequence. Sequence profiles and hidden Markov models built using homologous proteins ^6–10^ are often used to infer function by membership in a particular family, but these methods do not always identify specific functional residues and can misannotate proteins in mechanistically diverse families. ^11^ Additionally, structure and function are often conserved even when sequence similarity is very low, resulting in large numbers of false negatives for methods based on sequence alignment. ^12, 13^ Approaches based on identifying conserved sequence motifs within families can help to address these issues. ^14, 15^ However, these methods suffer from similar limitations as sequences diverge, resulting in high false positive and false negative rates, especially when the functional residues are far apart in sequence. ^16^ More generally, sequence-based methods cannot capture the complex 3D conformations and physicochemical interactions required to accurately define a functional site or inform opportunities to engineer or mutate specific residues.

Recently, methods have applied machine learning to predict function from sequence ^17, 18^ or structure. ^19^ However, like profile-based methods, these lack the local resolution necessary to identify specific functional sites, and their reliance on non-specific functional labels such as those provided by Gene Ontology terms ^20^ often limits practical utility. ^21^ Machine learning approaches that focus on local functional sites are either specific to a particular type of site (e.g. ligand binding ^22, 23^, enzyme active sites ^24^) or require building specific models for each functional site of interest, ^25, 26^ which can be computationally expensive and demands sufficient data to train an accurate model.

A major consideration for building generalizable machine learning models for protein sites is the choice of local structure representation. FEATURE, ^27^ a hand-crafted property-based representation, has shown utility for many functionally-relevant tasks. ^25, 28, 29^ However, FEATURE uses heterogeneous features (a mix of counts, binary, and continuous) which are more difficult to train on and meaningfully compare in high dimensions. Additionally, FEATURE consists of radial features without considering orientation and does not account for interactions between atoms in 3D, leading to loss of information. ^26^ Deep learning presents an attractive alternative by enabling the extraction of features directly from raw data, ^30^ but the high complexity of deep learning models means that they require large amounts of labeled data. To address this, a paradigm has emerged in which models are pre-trained on very large unlabeled datasets to extract robust and generalizable features which can then be “transferred” to downstream tasks. ^31, 32^ This approach has been successfully applied to learn representations of small molecules ^33, 34^ and protein sequences, ^17, 35, 36^ but there are few examples of representations learned directly from 3D structure. Initial efforts focus on entire proteins rather than sites and operate only at residue-level resolution. ^37, 38^

We address these issues by developing COLLAPSE (Compressed Latents Learned from Aligned Protein Structural Environments), a framework for functional site characterization, identification, and comparison which (1) focuses on local structural sites, defined as all atoms within a 10 Å radius of a specific residue; (2) captures complex 3D interactions at atom resolution; (3) works with arbitrary sites, regardless of the number of known examples; and (4) enables comparison between sites across proteins. COLLAPSE combines self-supervised methods from computer vision, ^39^ graph neural networks designed for protein structure, ^40, 41^ and multiple sequence alignments of homologous proteins to learn 512-dimensional protein site embeddings that capture structure-function relationships both within and between proteins.

Self-supervised representation learning refers to the procedure of training a model to extract high-level features from raw data using one or more “pretext tasks” defined using intrinsic characteristics of the input data. The choice of pretext task is critical to the utility of the learned representations. A popular class of methods involves minimizing the distance between the embeddings of two augmented versions of the same data point (for example, cropped and rotated views of the same image), thereby learning a representation that is robust to noise which is independent of the fundamental features of the original data. ^39, 42, 43^ Since function is largely conserved within a protein family, we draw an analogy between homologous proteins and augmented views of the same image. Specifically, we hypothesized that by pulling together the embeddings of corresponding sites in homologous proteins, we could train the model to learn features which capture the site’s structural and functional role. In this scheme, sequence alignments are used to identify correspondences between amino acids, which are then mapped to 3D structures to define the structural site surrounding each residue (Figure 1, Section 2.2).

**Figure 1.**
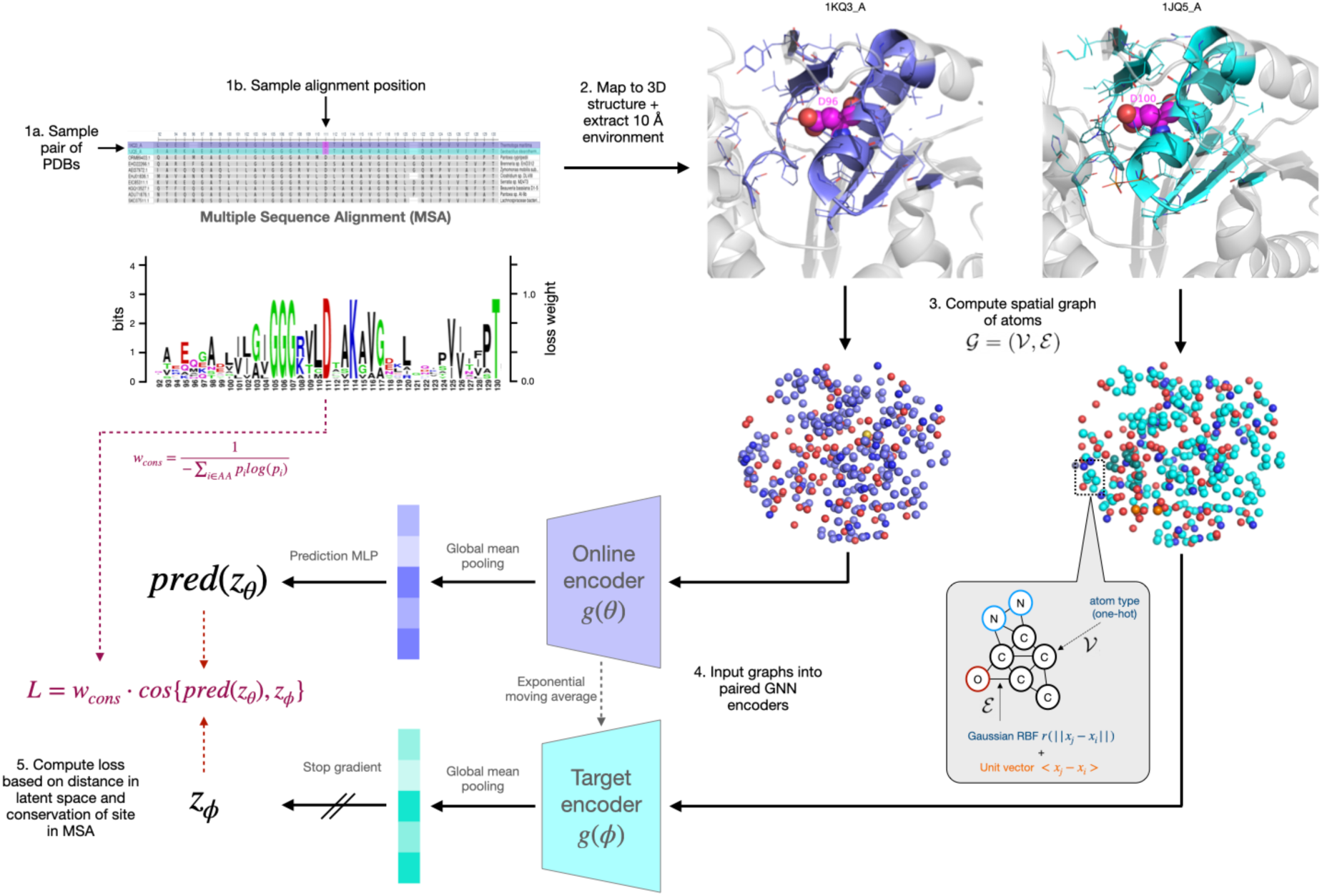
Schematic of a single iteration of COLLAPSE algorithm. Clockwise from the top left, we show **(1a,b)** the process of sampling a pair of sites from the MSA, **(2)** extracting the corresponding structural environments, and **(3)** converting into a spatial graph. The inset shows the node and edge featurization scheme. Finally, we show **(4)** a schematic of the network architecture, consisting of paired graph neural networks followed by mean pooling over all nodes to produce site embeddings. **(5)** These embeddings are then compared using a loss function weighted by the conservation of the position in the MSA, as shown by the sequence logo in center left.

Pre-trained representations are typically used in one of two settings: (1) transfer learning, which leverages general representations to improve performance on problem-specific supervised tasks where access to labeled data is limited; and (2) extracting insights about the underlying data from the learned embedding space directly (e.g. via visualization or embedding comparisons). ^44^ In this paper, we illustrate the utility of COLLAPSE protein site in both settings. First, we demonstrate that COLLAPSE generalizes in a transfer learning setting, achieving competitive or best-in-class results across a range of downstream tasks. Second, we describe two applications that demonstrate the power of our embeddings for protein function analysis without the need to train any downstream models: an iterated search procedure for identifying similar functional sites across large protein databases, and a method for efficiently annotating putative functional sites in an unlabeled protein. All datasets, models, functionality, and source code can be found in our Github repository (https://github.com/awfderry/COLLAPSE).

## 2. Results

### 2.1. Intrinsic evaluation of COLLAPSE embeddings

To evaluate the extent to which COLLAPSE embeddings capture relevant structural and functional features, we embedded the environments of all residues in a held-out set consisting of proteins with varying levels of sequence similarity to proteins in the training set. First, we find that the degree of similarity between embeddings of aligned sites is correlated with the level of conservation of that site in the multiple sequence alignment (MSA) (Figure 2a). Even at less than 30% conservation, aligned sites are significantly more similar on average than a randomly sampled background of non-aligned sites (*p* < 1 × 10^−15^).

**Figure 2.**
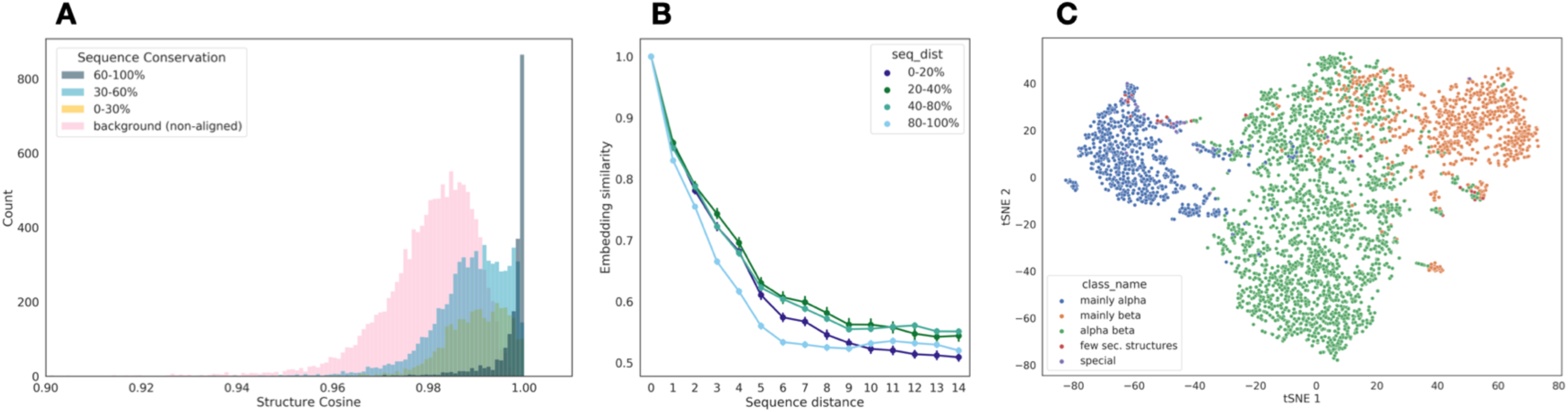
Analysis of learned embeddings. **(a)** Raw cosine similarity distributions (i.e. before quantile transformation) of aligned sites, binned by the sequence conservation of the corresponding column of the MSA. Highly conserved positions also have highly similar embeddings, but even less conserved positions have more similar embeddings than randomly sampled non-aligned sites (in pink). **(b)** Spatial sensitivity of embedding similarity, as measured by the sequence distance between two sites. Results are stratified by the average distance to the closest training protein, demonstrating that neighboring embeddings can be readily distinguished even for proteins with very low similarity to the training set. **(c)** tSNE projection of average protein-level embeddings for single-domain chains, colored by the highest-level CATH class, showing that embeddings effectively capture secondary structure patterns.

We also confirmed that our embeddings capture local information at a residue-level resolution, meaning that neighboring environments can be effectively distinguished from each other. Indeed, the normalized cosine similarity between residue embeddings decreases between the residues in sequence increases (Figure 2b). This effect generalizes even to proteins far away from the training set in sequence identity. Finally, among chains with a single fold according to CATH 4.2 ^45^ (*n* = 11,270), the top-level structural class can be distinguished clearly in protein-level embeddings, suggesting that secondary structure is a major feature captured by COLLAPSE (Figure 2c). Lower levels of the CATH hierarchy also cluster clearly in low-dimensional space (Figure S1).

### 2.2. Transfer learning and fine-tuning to improve performance on supervised tasks

To assess COLLAPSE in a transfer learning context, we use ATOM3D, a suite of benchmarking tasks and datasets for machine learning in structural biology. ^46^ We selected two tasks from ATOM3D which focus on protein sites: protein interface prediction (PIP) and mutation stability prediction (MSP). We compare performance to the ATOM3D reference models and to the task-specific GVP-GNN reported in Jing et al. (2021), ^41^ which is state-of-the-art for all tasks. Table 1 reports the results both with and without fine-tuning the embedding model parameters. Without fine-tuning, COLLAPSE embeddings and a simple classifier achieve results comparable or better than the ATOM3D reference models trained specifically for each task. Fine-tuning improves performance further, achieving state-of-the-art on PIP and comparable performance to the GVP-GNN on MSP, outperforming FEATURE as well as the ATOM3D baselines (Table S6). As an external evaluation, we also evaluated COLLAPSE on the prediction of protein-protein interaction sites compared to MaSIF, a deep learning model designed for protein surfaces ^47^. Our models perform comparably to MaSIF and better than baseline methods despite training on orders of magnitude fewer sites on the protein surface (Figure S8).

**Table 1.**
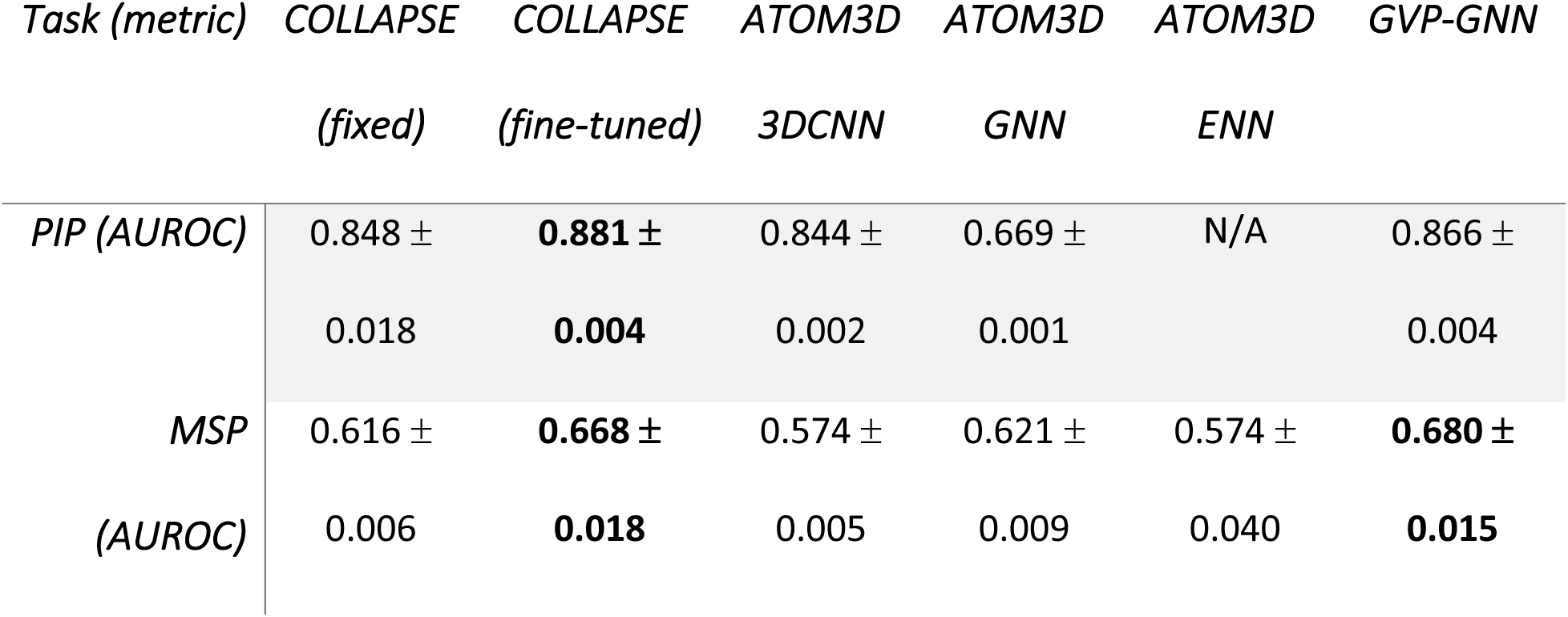
Performance of models trained on ATOM3D benchmark tasks. Comparisons are made with ATOM3D reference architectures—3D convolutional neural network (3DCNN), graph neural network (GNN), and equivariant neural network (ENN)—as well as the geometric vector perceptron (GVP-GNN) results reported in Jing et al. (2021) ^41^, which is state-of-the-art for these datasets. The metric is area under the receiver operator characteristic curve (AUROC), and we report mean and standard deviation across three training runs. Numbers in bold indicate best performance on each task (within one standard deviation).

| *Task (metric)* | *COLLAPSE* | *COLLAPSE* | *ATOM3D* | *ATOM3D* | *ATOM3D* | *GVP-GNN* |
| --- | --- | --- | --- | --- | --- | --- |
|  | *(fixed)* | *(fine-tuned)* | *3DCNN* | *GNN* | *ENN* |  |
| *PIP (AUROC)* | 0.848 ± | **0.881 ±** | 0.844 ± | 0.669 ± | N/A | 0.866 ± |
|  | 0.018 | **0.004** | 0.002 | 0.001 |  | 0.004 |
| *MSP* | 0.616 ± | **0.668 ±** | 0.574 ± | 0.621 ± | 0.574 ± | **0.680 ±** |
| *(AUROC)* | 0.006 | **0.018** | 0.005 | 0.009 | 0.040 | **0.015** |

### 2.3. Building functional site prediction models using COLLAPSE embeddings

We train prediction models for 10 functional sites defined by the Prosite database, ^14^ which identifies local sites using curated sequence motifs. On sites labeled true positive (TP) by PROSITE, COLLAPSE outperforms the analogous FEATURE models and perform comparably or better than task-specific 3DCNN models trained end-to-end, achieving greater than 86% recall on all sites at a threshold of 99% precision. Prosite also provides false negatives (FNs; true proteins which are not recognized by the Prosite pattern) and false positives (FPs; proteins which match the Prosite pattern but are not members of the functional family). Table 2 shows the number of proteins correctly reclassified by the models trained on TP sites. For all families, COLLAPSE correctly identifies a greater or equal number of FN proteins compared to FEATURE and 3DCNN classifiers. The improvement is notable in some cases, such as a 162.5% increase in proteins detected for IG_MHC, a 37.5% increase for ADH_SHORT, and a 17.6% increase for EF_HAND_1. For four of the seven proteins with FP data, we correctly rule out all FPs. For ADH_SHORT and EF_HAND_1, we perform 9.1% and 4.0% worse relative to 3DCNN, respectively, but this slight increase in FPs is not substantial relative to the improvement in FNs recovered for these families. To evaluate our embeddings beyond PROSITE, we also trained functional residue prediction models on four benchmark datasets from Xin et al. (2011). ^48^ Models based on COLLAPSE embeddings perform significantly better than the best-performing method on the prediction of zinc binding sites and enzyme catalytic sites (15.3% and 9.3% improvement, respectively), and within 10% of the best-performing method on DNA binding and phosphorylation sites (Table S5).

**Table 2.**
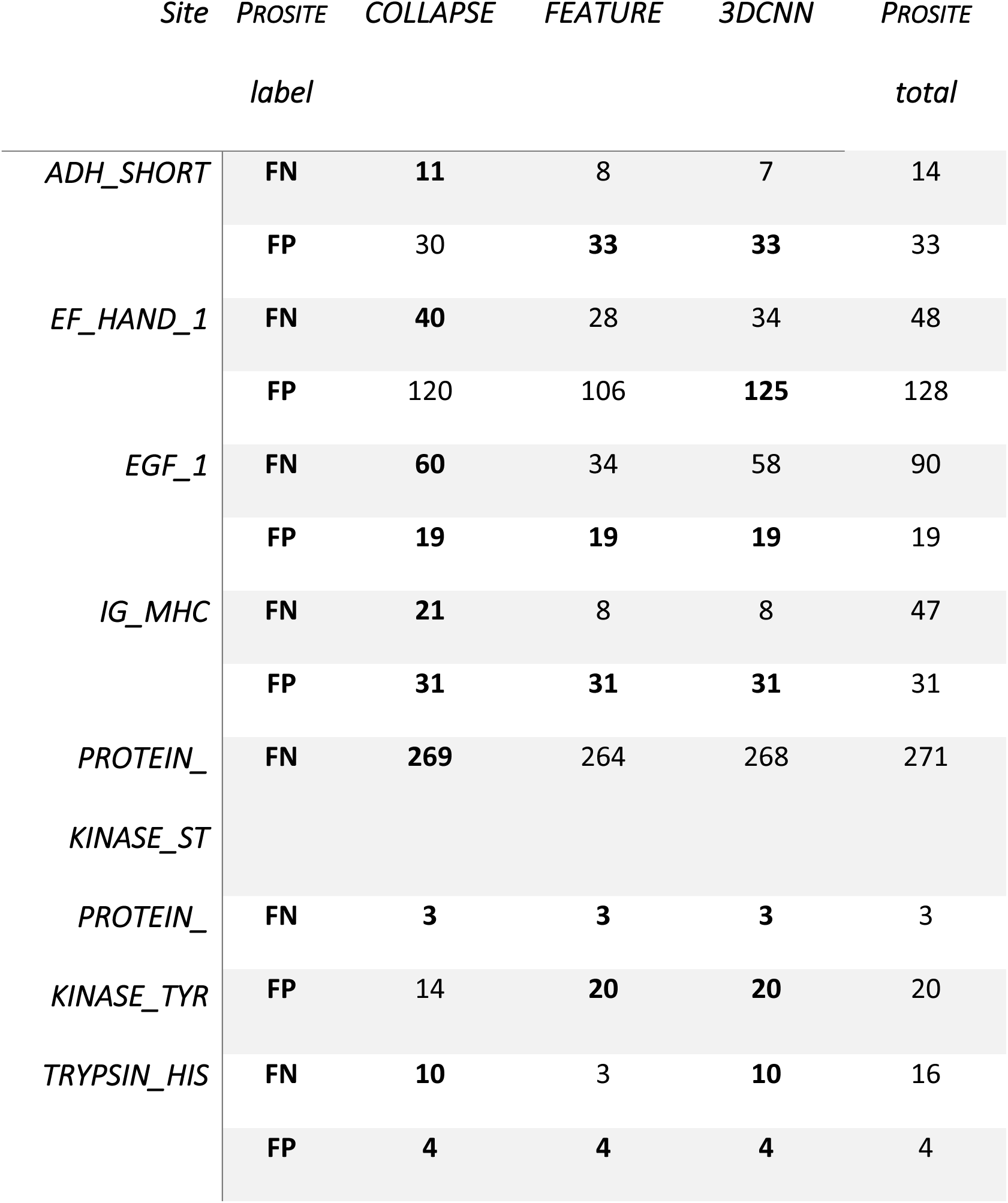
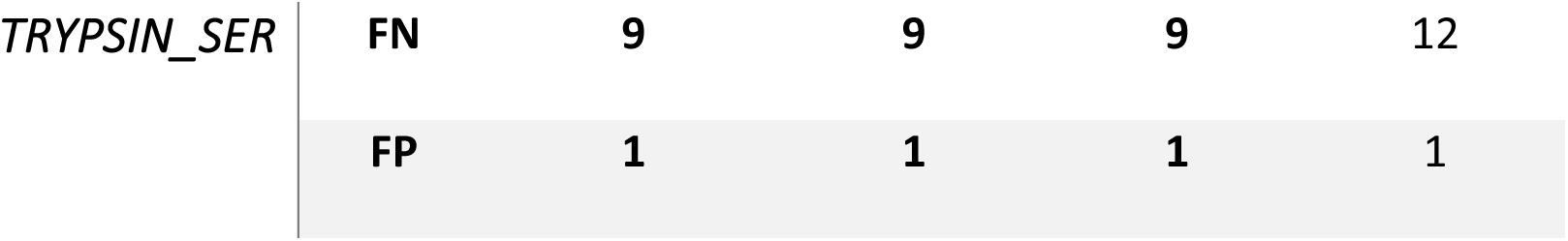
Performance of models trained on Prosite TP/TN on held-out Prosite FP/FN annotations. Comparisons are made with FEATURE and 3DCNN numbers as reported in Torng et al. (2019) ^26^. The number of proteins which are correctly reclassified (i.e. FPs predicted as negative, FNs predicted as positive) is reported for each method (higher is better). Numbers in bold indicate best performance on each site.

### 2.4. Iterative search for functional sites across protein databases

While COLLAPSE embeddings can be used to train highly accurate models for functional site detection, we can only train such models for those functional sites for which we have sufficient training examples. Another way to understand the possible function of a site is to analyze similar sites retrieved from a structure database. The set of hits retrieved by this search may contain known functional annotations or other information which sheds light on the query site. We use iterative COLLAPSE embedding comparisons to perform such a search across the PDB. We investigate the performance of this method on the Prosite dataset while varying two parameters: the number of iterations and the p-value cutoff for inclusion at each iteration. The method generally achieves high recall and precision after 2–5 iterations at a p-value cutoff of 5 × 10^−3^ to 5 × 10^−4^ (Figure 4; Figure S4). Notably, when evaluating on the FN and FP subsets, our search method even outperforms the cross-validated models on some sites (e.g. IG_MHC, Figure 4a). However, the precision and recall characteristics vary widely across families; in some cases it predicts the same set of proteins as the trained model (e.g. TRYPSIN_HIS; Figure 4b), while in others it performs worse (e.g. EF_HAND_1; Figure 4c). Importantly, the method requires no training and is very efficient: runtime per iteration scales linearly with the size of the query set and with database size (Figure S5).

**Figure 3.**
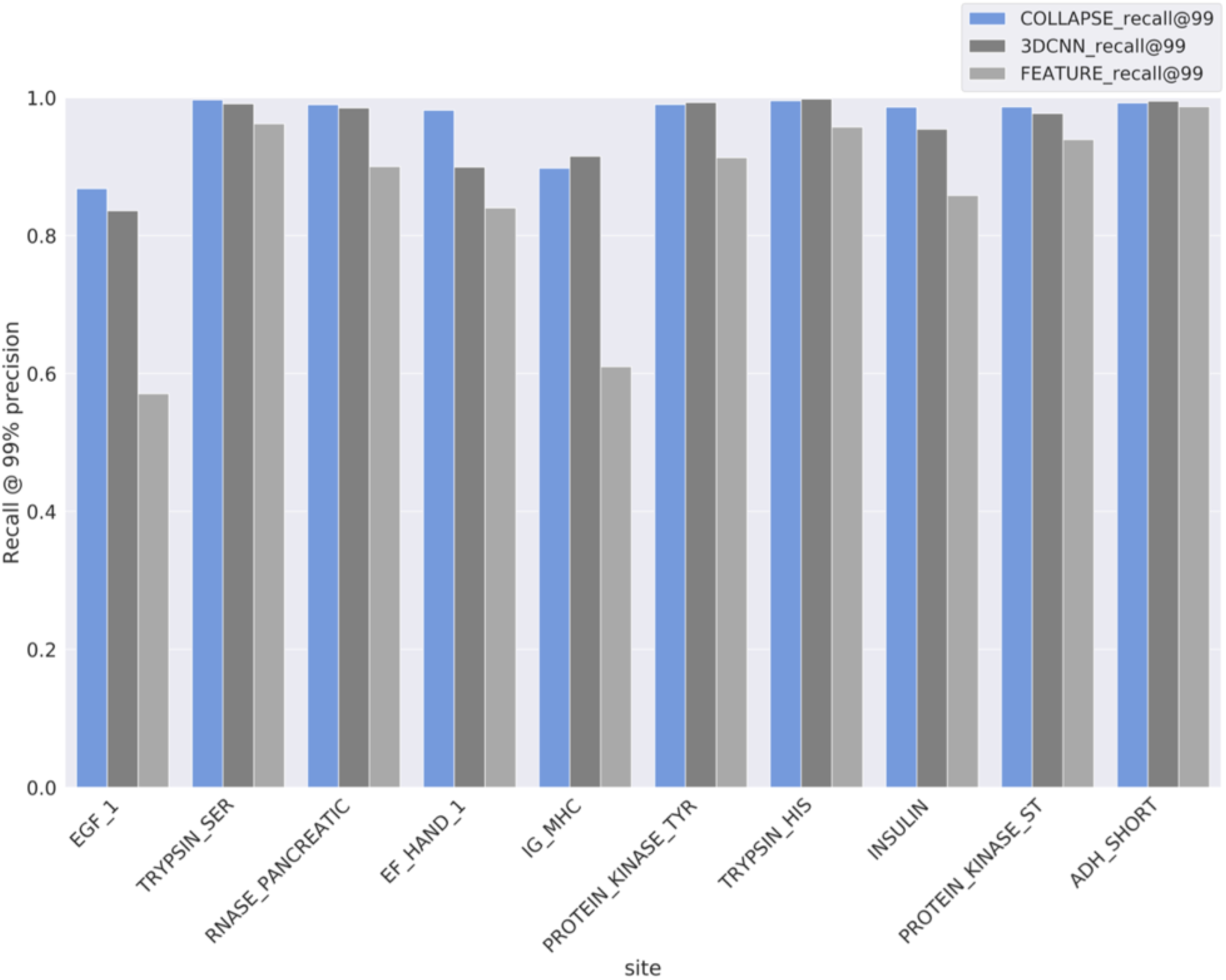
Performance of models trained on true positives from 10 Prosite functional sites in 5-fold cross-validation: COLLAPSE embeddings + SVM (blue), 3DCNN trained end-to-end (dark gray), and FEATURE vectors + SVM (light gray). Metric is the recall for all TP annotations at a threshold which produces 99% precision. COLLAPSE achieves better recall than FEATURE and better or comparable recall to the 3DCNN.

**Figure 4.**
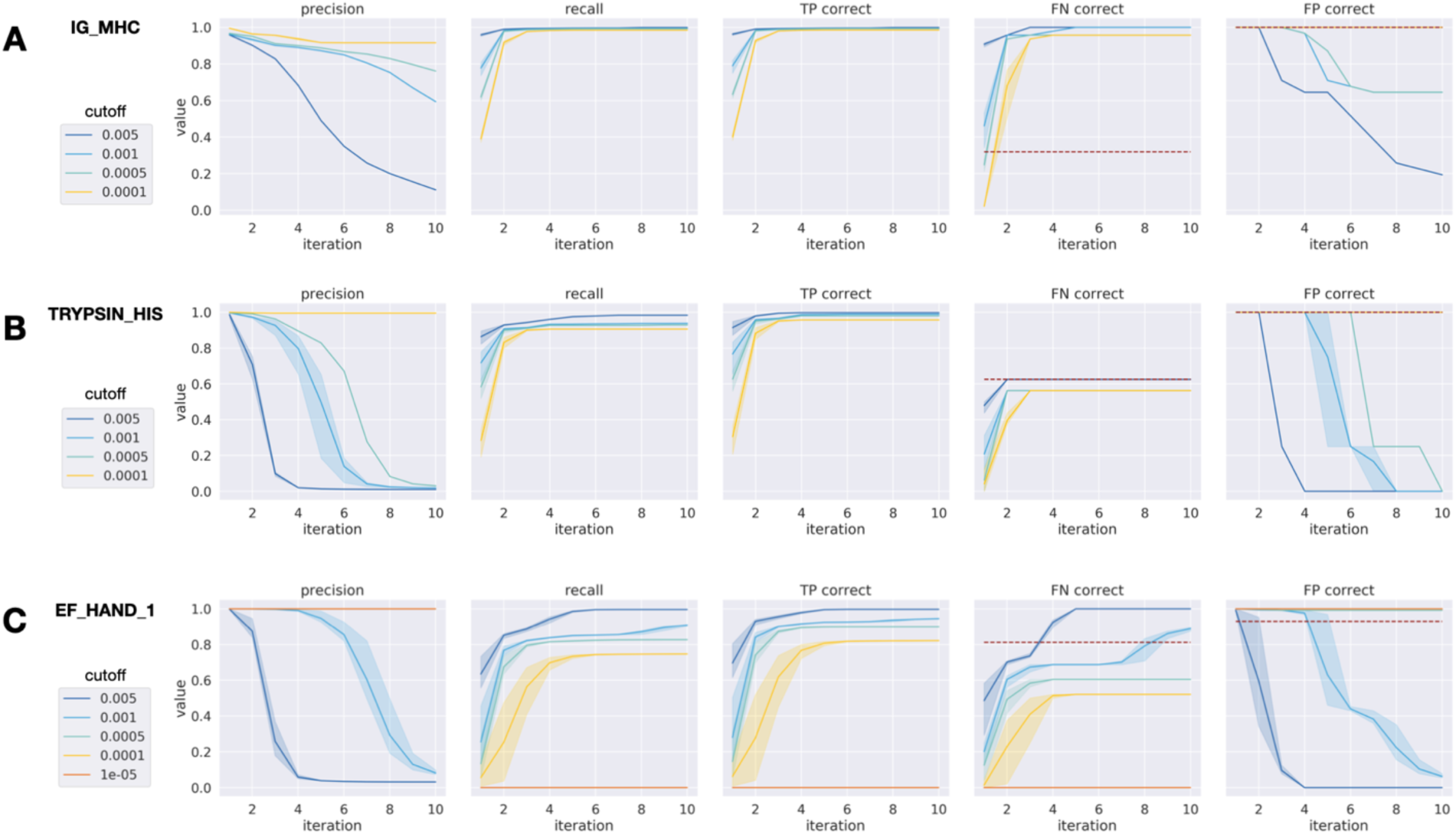
Iterated functional site search performance per iteration for three Prosite families. Colors denote different user-specified empirical p-value cutoffs and error bars represent variance over three randomly sampled queries. From left to right, metrics shown are: precision across all results (including TP, FP, and FN), recall across all results, proportion of TP sites predicted correctly, proportion of FN sites predicted correctly, and proportion of FP sites predicted correctly. For FN and FP, the performance of our CV-trained models is shown as a red dashed line. Sample error is shown for three random starting queries. The three families shown are **(a)** IG_MHC, **(b)** TRYPSIN_HIS, and **(c)** EF_HAND_1, in order of relative performance compared to CV-trained models. While the performance characteristics vary across sites, the number of iterations and p-value cutoff can be tuned to achieve good performance.

### 2.5. Annotation of functional sites in protein structures

Our iterative search method assumes that a site of interest has already been identified. However, when a new protein is discovered and its structure is solved, the locations of functional sites are often unknown. By comparing local environments in the protein’s structure to those contained in databases of known functional sites, we can predict which sites are likely to be functional. Figure 5 shows two example annotations using a modified mutual best hit criterion against a reference database consisting of embeddings from Prosite and the Catalytic Site Atlas (CSA). First, we show the structure of meizothrombin, a precursor to thrombin and a trypsin-like serine protease with a canonical His-Asp-Ser catalytic triad. Our method correctly identifies all three residues as belonging to the trypsin-like serine protease family in Prosite (Figure 5a). Hits against the CSA, which are more specific, also include closely homologous proteins such as C3/C5 convertase. The associated kringle domain is also identified by its characteristic disulfide bond. Second, we show the structure of beta-glucuronidase (Figure 5b), a validation set protein which has no homologs in the training set. We correctly identify all four catalytic residues defined by the CSA (in yellow), as well as Prosite signatures corresponding to the glycosyl hydrolases family 2, the family which contains beta-glucuronidase.

**Figure 5.**
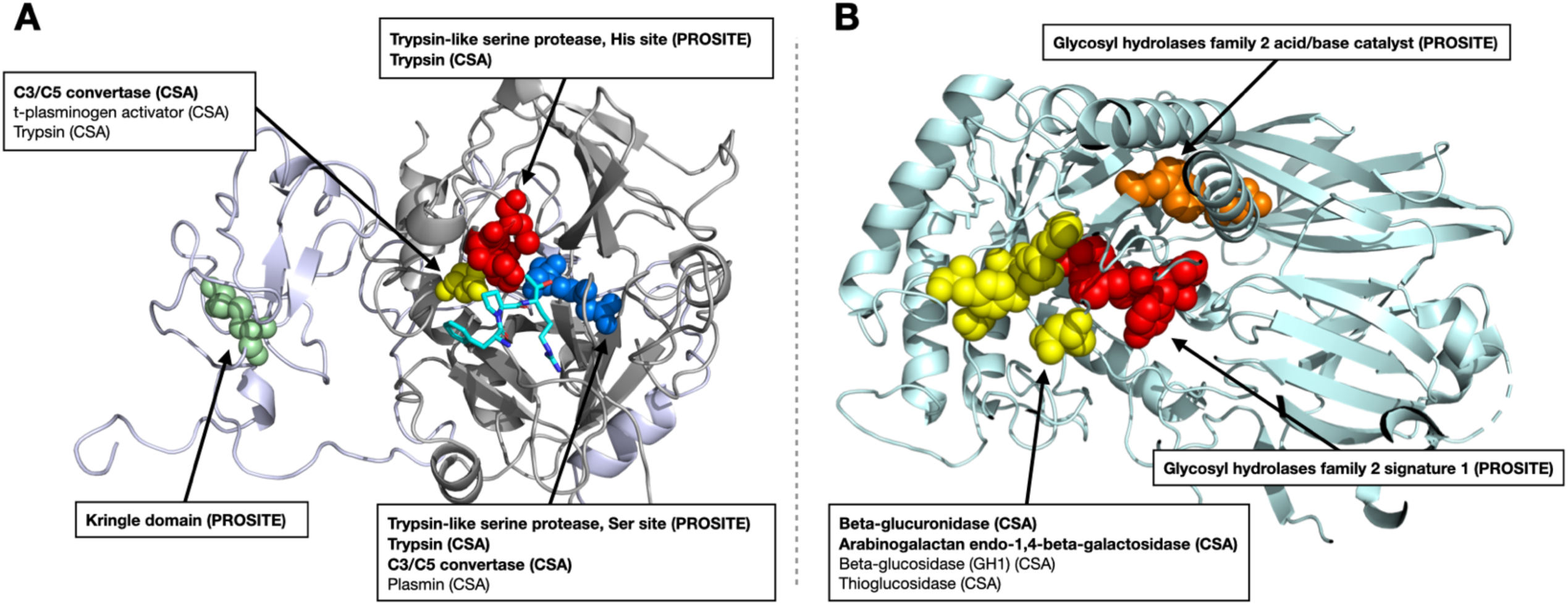
Results of functional annotation tool applied to **(a)** meizothrombin (PDB ID 1A0H) and **(b)** beta-glucuronidase (PDB ID 3HN3), both at *p* < 1 × 10^−4^. No member of the beta-glucuronidase family is in the training set (maximum sequence identity 2.8%). Functional residues identified by our method are shown as spheres, with colors corresponding to the functional site. Hits labeled in bold are also significant at a more stringent cutoff (*p* < 5 × 10^−5^). All hits represent either the correct function or those of very closely related proteins, showing that COLLAPSE is effective for annotation of proteins whether or not similar proteins are present in the training set.

## 3. Discussion

### 3.1. The COLLAPSE self-supervised training framework enables the learning of rich, informative embeddings of functional sites

The utility of COLLAPSE embeddings for functional analysis derives from several key features of the training algorithm. First, the use of homology as a source of self-supervision signal allows the model to learn patterns of structural conservation across proteins, imbuing the model with a biological inductive bias towards features that may be important to the protein’s function. Such patterns could in theory be learned by a model which sees each protein independently, but it would require much more data and training time to identify subtle signals across disparate proteins. The bootstrap training objective not only forces the model to learn a distance function for comparing embeddings that meaningfully captures the functional relationship between sites, but by sampling pairs of proteins and residues each iteration it also greatly increases the effective size of the training set relative to models that take in a single protein (or even residue) at a time. Indeed, we have found empirically that non-bootstrap objectives (e.g. those based on masked residue modeling or autoencoders) produce representations that are much less informative for functional tasks. While MSAs have proved crucial to the success of sequence-based models, to our knowledge this is the first time they have been used to provide a supervision signal for a structure-based model.

Second, by focusing on local protein sites, our embeddings are more precise and flexible than models which produce a single representation of an entire protein. COLLAPSE embeddings can be used for arbitrary tasks on the level of single residues or even individual functional atoms, to detect important regions in proteins, and to identify functional relationships between proteins even if they are divergent in sequence or global fold. Moreover, by aggregating over multiple residues or entire proteins, site-specific embeddings can also be applied to domain-level or full-protein tasks. For simple tasks such as distinguishing high-level CATH class (Figure 2c), a naive approach of averaging over all residues is sufficient, but more complex aggregation methods will be necessary for many problems. As a proof of concept, we trained a simple recurrent neural network (RNN) model on sequences of COLLAPSE embeddings to make protein-level predictions of fold and enzyme class (Supplementary Note 9). Compared to 16 other sequence and structure based methods, our RNNs trained on COLLAPSE embeddings rank in the top five across all tasks and testing sets (Figure S7).

Finally, by using an atomic graph representation and a GVP-GNN encoder, COLLAPSE captures all inter-atomic interactions (in contrast to methods which operate at a residue level) and produces representations that are fully equivariant to 3D rotation and translation. The importance of capturing local structural features in functional site analysis is demonstrated by the improved performance of COLLAPSE relative to sequence-based methods such as MMSeqs2 (Table S4), especially on the more difficult FN and FP proteins. While sequence embeddings from ESM-1b ^50^ do produce very good predictive models for certain sites, they dramatically overfit to others (notably IG_MHC) and generally underperform across all sites in search and annotation applications (Figures S9–S10, Supplementary Note 5). This suggests that sequence and structure representations are complementary, and many tasks may benefit from a combination of the two approaches.

### 3.2. COLLAPSE produces general-purpose representations which facilitate the improvement of computational methods for diverse applications in protein function analysis

COLLAPSE is effective in transfer learning and as fixed embeddings, and generalizes across tasks that require the model to learn different aspects of the protein structure-function relationship. Although it may not be the state-of-the-art on every task, its competitive performance in every context we tested demonstrates its utility as a general-purpose representation. This flexibility makes COLLAPSE embeddings ideal for not only building new predictive models and performing comparative analyses, but also for easily incorporating structural information into existing computational methods. Given the improved performance over FEATURE across our benchmarks, we also expect that substituting COLLAPSE embeddings will lead to improved performance for most applications addressed by the FEATURE suite of methods ^28, 29, 51^.

Another important aspect which sets COLLAPSE apart from task-specific machine learning methods is the ability to perform meaningful comparisons between functional sites directly in the embedding space. Due to the bootstrap pre-training objective, the embedding distance provides a functionally relevant distance measure for comparing sites. Note that for all comparisons, our method of standardizing embedding comparisons is critical for determining their statistical significance as well as increasing their effective range (Supplementary Note 1). We demonstrate the benefits of using comparisons directly in the embedding space by developing methods for functional site search and annotation, both of which are efficient, generalizable, and allow a user to tune the sensitivity and specificity of the results. For example, for discovery applications it may be desirable to optimize for sensitivity at the cost of more false positives, while prioritizing drug targets for experimental validation may require greater specificity.

The ability of iterative nearest-neighbor searches in the embedding space to identify known sites in Prosite demonstrates that functional sites cluster meaningfully in the embedding space. The effect of changing input parameters (number of iterations and p-value cutoff) on the sensitivity and specificity of the results varies somewhat across functional families. In some cases (notably IG_MHC), this method achieves better sensitivity for FNs than even machine learning models trained using CV, while in others (EF_HAND_1, PROTEIN_KINASE_TYR) it cannot achieve this without a significant drop in precision. This is likely due to differences in structural conservation between sites, whereby sites which are more structurally heterogeneous are more difficult to fully capture using a nearest-neighbor approach than a trained model which can learn to recognize diverse structural patterns. However, since training an accurate model requires access to a representative training dataset which is not always available, we consider our search method to be a powerful complement to site-specific models in cases where labeled data is scarce or where the similarity to a specific query is important. We also note that while structural search methods exist for full proteins ^52, 53^ or binding sites, ^29, 54, 55^ ours is the first search tool specifically designed for arbitrary local structural sites.

Functional annotation of novel protein structures is of great value to the structural biology and biochemistry communities, but there are few tools for doing so at the residue level. COLLAPSE provides a method for residue-level annotation which is efficient and tunable, making it suitable for both screening and discovery purposes. As shown by the examples in Figure 5, the method identifies known functional annotations while limiting false positives to closely related homologs, even when the input is not related to any protein in the training set (<5% sequence identity for beta-glucuronidase). Confidence in a new prediction’s accuracy can be assessed by its significance level or more sophisticated evaluations for example, multiple neighboring residues being annotated with the same function may imply a more likely correct prediction. Additionally, because we can identify many sites on a single protein, it is also possible to identify previously unknown alternative functional sites or allosteric sites. Importantly, all predictions can be explained and cross-referenced by rich metadata from the reference data sources, enhancing trust and usability. Of the PDBs returned for true positive sites in meizothrombin and beta-glucuronidase, 45.5% (20/44) and 87.5% (14/16), respectively, were not hits in a protein PSI-BLAST search with standard parameters, demonstrating the value of local structural comparisons for functional annotation. Additionally, the method is easy to update and extend over time via the addition of new sources of functional data, and reference databases can even be added or removed on a case-by-case basis.

### 3.3. Advances in protein structure prediction provide great opportunities for expanding functional analysis and discovery

COLLAPSE depends on the availability of solved 3D protein structures in the PDB. This restricts not only the number of homologous proteins that can be compared at each training step, but also the set of protein families which can even be considered—less than one third of alignments in the CDD contained at least two proteins with structures in the PDB. Including structures from AlphaFold Structure Database ^56^ would dramatically increase the coverage of our training dataset, but the utility of including predicted structures alongside experimentally solved structures in training or evaluation of machine learning models still needs to be evaluated. ^57^ A preliminary evaluation of our annotation method on the predicted structure for meizothrombin reveals high agreement with the corresponding PDB structure (Figure S6) despite a root-mean-square deviation of 3.67 Å between the two structures, suggesting that COLLAPSE may already generalize to AlphaFold predictions for some proteins. Given this finding, we anticipate that COLLAPSE will be a powerful tool for functional discovery within the AlphaFold database, which has already yielded several novel insights ^58, 59^.

In summary, COLLAPSE is a general-purpose protein structure embedding method for functional site analysis. We provide a Python package and command-line tools for generating embeddings for any protein site, conducting functional site searches, and annotating input protein structures. We also provide downloadable databases of embeddings for a non-redundant subset of the PDB and for known functional sites. We anticipate that as more data becomes available, these tools will serve as a catalyst for data-driven biological discovery and become a critical component of the protein research toolkit.

## 4. Materials and Methods

### 4.1. Training dataset and data processing

COLLAPSE pre-training relies on a source of high-quality protein families associated with known structures and functions, as well as multiple sequence alignments (MSAs) in order to define site correspondences. We use the NCBI-curated subset of the Conserved Domain Database (CDD), ^60, 61^ which explicitly validates domain boundaries using 3D structural information. We downloaded all curated MSAs from the CDD (n=17,906 as of Sep. 2021) and filtered out those that contained less than two proteins with structures deposited in the PDB. After removing chains with incomplete data or which could not be processed properly, this resulted in 5,643 alignments for training, corresponding to 16,931 PDB chains (Figure S2). We then aligned the sequences extracted from the ATOM records in each PDB chain to its MSA, without altering the original alignment, thus establishing the correct mapping from alignment position to PDB residue number. As a held-out set for validation, we select 1,370 families defined by PFAM ^6^ which do not share a common superfamily cluster (as defined by the CDD) with any training family. We then bin these families based on the average sequence identity to the nearest protein in the training dataset and sample five families from each bin, resulting in 50 validation families with varying levels of similarity to the training data (Table S1).

#### 4.1.1. Definition of sites and environments

In general, we define protein sites relative to the location of the relevant residues. Specifically, we define the environment surrounding a protein site as all atoms within 10 Å radius of the functional center of the central residue. The functional center is defined as the centroid of the functional atoms of the side chain as defined by previous work. ^26, 27^ For residues with two functional centers (Trp and Tyr), during training one is randomly chosen at each iteration, and at inference time the choice depends on the specific application (i.e. if the function being evaluated depends on the aromatic or polar group; see Table S2). If the functional atom is not known (e.g. for annotating unlabeled proteins), we take the average over all heavy side-chain atoms.

#### 4.1.2. Empirical background calculation

To make comparisons more meaningful and to provide a mechanism for calculating statistical significance, we quantile-transform all cosine similarities relative to an empirical cosine similarity distribution. To compute background distributions, we use a high-resolution (<2.0 Å), non-redundant subset of the PDB at 30% sequence identity provided by the PISCES server ^62^ (5,833 proteins). We compute the embeddings of 100 sites from each structure, corresponding to five for each amino acid type, sampled with replacement. Exhaustively computing all pairwise similarities is computationally infeasible, so we sample *n* = 50,000 pairs of environments and compute the cosine similarity of each. We performed this procedure to generate empirical similarity distributions (*S*_1_, …, *S_n_*) for the entire dataset and for each amino acid individually (Figure S3). Cosine similarities (*s*) are then quantile-transformed relative to the relevant empirical cumulative distribution function:

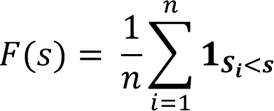

The p-value for any embedding comparison is then defined as 1– *F*(*s*), or the probability that a randomly sampled pair of embeddings is at least as similar as the pair in question. Amino acid– specific empirical backgrounds are used for functional site search and are aggregated into a single combined distribution for annotation. For the functional site–specific background used to filter hits during annotation, we use an empirical background computed by comparing each functional site embedding to the embeddings of the corresponding amino acid in the 30% non-redundant PDB subset.

### 4.2. COLLAPSE training algorithm

Each iteration of the COLLAPSE pre-training algorithm consists of the following steps, as shown in Figure 1. We trained our final model using the Adam optimizer ^63^ with a learning rate of 1e-4 and a batch size of 48 pairs for 1,200 epochs on a single TESLA V100 GPU. Model selection and hyperparameter tuning was evaluated using intrinsic embedding characteristics (Section 2.1) and ATOM3D validation set performance (Section 4.3). See Supplementary Note 2 for further discussion of hyperparameter selection and modeling choices.

*Step 1.* Randomly sample one pair of proteins from the MSA and one aligned position from each protein (i.e. there is not a gap in either protein). Map MSA column position to PDB residue number using the pre-computed alignment described in Section 2.2. Note that this step ensures that each epoch, a different pair of residues is sampled from each CDD family, effectively increasing the size of the training dataset by many orders of magnitude relative to a strategy which trains on individual proteins or MSAs.

*Step 2.* Extract 3D environment around each selected residue (Section 4.1.1). Only atoms from the same chain are considered. Waters and hydrogens are excluded but ligands, metal ions, and cofactors are included.

*Step 3.* Convert each environment into a spatial graph 𝒢 = (𝒱, ℰ). Each node in the graph represents an atom and is featurized by a one-hot encoding of the atom type 𝒱 ∈ { carbon (C), nitrogen (N), oxygen (O), fluorine (F), sulfur (S), chlorine (Cl), phosphorus (P), selenium (Se), iron (Fe), zinc (Zn), calcium (Ca), magnesium (Mg), and “other” }, representing the most common elements found in the PDB. Edges in the graph are defined between any pair of atoms separated by than 4.5 Å. Following Jing et al. (2021), ^41^ edges between atoms (*i*, *j*) with coordinates (*x_i_*, *x_j_*) are featurized using (1) a 16-dimensional Gaussian radial basis function encoding of distance *r*(||*x_j_* – *x_i_*||) and (2) a unit vector < *x_j_* – *x_i_* > encoding orientation.

*Step 4.* Compute embeddings of each site. We embed each pair of structural graphs (𝒢_1_, 𝒢_2_) using a pair of graph neural networks, each composed of three layers of Geometric Vector Perceptrons (GVPs), ^40, 41^ which learn rotationally-equivariant representations of each atom and have proved to be state-of-the-art in a variety of tasks involving protein structure. ^41, 64^ We adopt all network hyperparameters (e.g. number of hidden dimensions) from Jing et al. (2021). ^41^ Formally, each GVP learns a transformation of the input graph into 512-dimensional embeddings of each node:

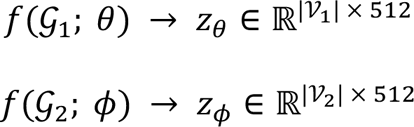

The final embedding of the entire graph is then computed by global mean pooling over the embeddings of each atom. While in principle, the two networks could be direct copies of each other (i.e. have tied parameters *θ* = *ϕ*), we adopt the approach proposed by Grill et al (2020) ^39^ which refers to the two networks as the *online encoder* and the *target encoder*, respectively. Only the online network parameters *θ* are updated by gradient descent, while the target network parameters *ϕ* are updated as an exponential moving average of *θ*:

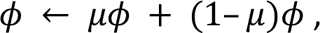

where *μ* is a momentum parameter which we set equal to 0.99. No gradients are propagated back through the target network, so only *θ* is updated based on the data during training. Intuitively, the target network produces a regression target based on a “decayed” representation, while the online network is trained to continually improve this representation. Only the online network is used to generate embeddings for all downstream applications.

*Step 5.* Compute loss and update parameters. The loss function is defined directly in the embedding space using the cosine similarity between the target network embedding *z*_*ϕ*_ ∈ ℝ^512^. and the online network embedding *z*_*θ*_ ∈ ℝ^512^. projected through a simple predictor network *pred*(*z*_*θ*_) ∈ ℝ^512^. This predictor network learns to optimally match the outputs of the online and target networks and is crucial to avoiding collapsed representations. To increase the signal-to-noise ratio and encourage the model to learn functionally relevant information, we weight the loss at each iteration by the sequence conservation *w_cons_* of that column in the original MSA (defined by the inverse of the Shannon’s entropy of amino acids at that position, ignoring gaps). To reduce bias, we include all proteins in the alignment curated by CDD for computing conservation, even those without corresponding structures. As a result of this, the loss function is expressed as:

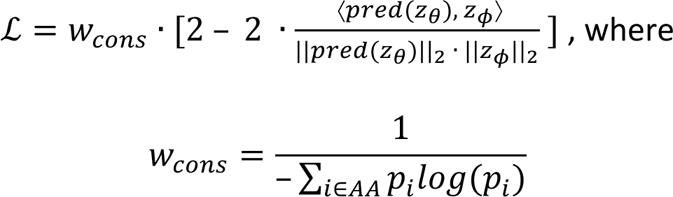

Finally, we symmetrize the loss by passing each site in the input pair through both online and target networks and summing the loss from each. This symmetrized loss is then used to optimize the parameters of the online network using gradient descent.

### 4.3. Transfer learning and fine-tuning

We retrieved the pre-processed PIP and MSP datasets from Townshend et al., ^46^ as well as the performance metrics for baseline models. Both datasets consist of paired residue environments, so we embed the environments surrounding each residue in the pair, concatenate the embeddings, and train a two-layer feed-forward neural network to predict the binary outcome (see Supplementary Note 3 for details). For the comparison with MaSIF, we obtained the MaSIF-site training and testing datasets from Gainza et al. ^47^ Since MaSIF operates on a dense mesh of points along the protein surface, we mapped all points to the nearest residue and embedded that residue’s local environment. We then added a two-layer feed-forward network to predict whether or not the residue is part of a protein-protein interaction site (Supplementary Note 4). For all tasks, we trained models with and without allowing the parameters of the embedding model to change (fine-tuned and fixed, respectively). Hyperparameters were selected by monitoring the relevant metric on the validation set.

### 4.4. Training site-specific models on Prosite data

We choose 10 sites presented in Torng et al. (2019), ^26^ selected because they are the most challenging to predict using FEATURE-based approaches. ^25, 26^ For each functional site, we train a binary classifier on fixed COLLAPSE embeddings in five-fold nested cross-validation (CV). The classifiers are support vector machines (SVMs) with radial basis function kernels and weighted by class frequency. Within each training fold, the inner CV is used to select the regularization hyperparameter *C* ∈ {0.1, 1, 10, 100, 1000, 5000} and the outer CV is used for model evaluation. To enable more accurate comparisons, we use the same dataset, evaluation procedures as Torng et al. (2019). ^26^ We benchmark against reported results for SVMs trained on FEATURE vectors (a direct comparison to our procedure) and 3D convolutional neural networks (3DCNNs) trained end-to-end on the functional site structures (the current state of the art for this task). We use Prosite FN/FP sites as an independent validation of our trained models, using an ensemble of the models trained on each CV fold and the classification threshold determined above. A site is considered positive if the probability estimate from any of the five fold models is greater than the threshold. Some proteins contain more than one site; in these cases, the protein is considered to be positive if any sites are predicted to be positive.

### 4.5. Iterated functional site search

First, we embed the database to be searched against using the pre-trained COLLAPSE model. For the results presented in Section 2.4, we use the same Prosite dataset used to train our cross-validated models to enable accurate comparisons. However, we also provide an embedding dataset for the entire PDB and scripts for generating databases for any set of protein structures. Then, we index the embedding database using FAISS, ^65^ which enables efficient similarity searches for high-dimensional data. For each site, we then perform the following procedure five times with different random seeds in order to assess the variability of results under different query sites. The input parameters are the number of iterations *n_iter_* and the p-value cutoff for selecting sites at each iteration *p_cutoff_*.

1. Sample a single site from the Prosite TP dataset (to simulate querying a known functional site), generate COLLAPSE embedding, and add to query set.
2. Compute effective cosine similarity cutoff *s_cutoff_* using the (1 – *p_cutoff_*) quantile of the empirical background for the functional amino acid of the query site (e.g. cysteine for an EGF_1 site).
3. Compare embedding(s) of query to database and retrieve all neighbors within *s*_4CB5DD_of the query.
4. Add all neighbors to query set and repeat Step 3 *n_iter_* times. Note that when there is more than one query point, neighbors to *any* point in the query are returned.
5. Compute precision and recall of final query set, using Prosite data as ground truth.

### 4.6. Protein site annotation

Instead of a database of all protein sites, the annotation method requires a database of known functional sites. We use all true positive sites defined in PROSITE. For each pattern, we identify all matching PDBs using the ScanProsite tool ^14^ and extract the residues corresponding to all fully conserved positions in the pattern (i.e. where only one residue is allowed). The environment around each residue is embedded using COLLAPSE. We also embed all residues in the Catalytic Site Atlas (CSA), a curated dataset of catalytic residues responsible for an enzyme’s function. All data processing matches the pre-training procedure. The final dataset consists of 25,407 embeddings representing 1,870 unique functional sites.

The annotation method operates in a similar fashion to the search method, where each residue in the input protein is embedded and compared to the functional site database. Any residue that has a hit with a p-value below the pre-specified cutoff is returned as a potential functional site. To filter out false positives due to common or non-specific features (e.g. small polar residues in alpha-helices), we also remove hits which are not significant against the empirical distribution specific to that functional site (Section 4.1.2). This results in a modified mutual best hit criterion with two user-specified parameters: the residue-level and site-level significance thresholds. Along with each hit is the metadata associated with the corresponding database entry (PDB ID, functional site description, etc.) so each result can be examined in more detail. For the examples presented we remove all ligand atoms from the input structure to reduce the influence of non-protein atoms on the embeddings.

## 5 Acknowledgments

We thank Kristy Carpenter, Delaney Smith, Adam Lavertu, and Wen Torng for useful discussions. Computing for this project was performed on the Sherlock cluster; we would like to thank Stanford University and the Stanford Research Computing Center for providing computational resources and support. A.D. is supported by LM012409 and R.B.A. is supported by NIH GM102365 and Chan Zuckerberg Biohub.

## Supplementary Materials

### Supplementary Note 1

The range of cosine similarities is relatively small (∼0.9–1.0), which we attribute to the use of mean pooling over atoms in the encoder’s graph aggregation step. A different choice of aggregation function may produce larger dynamic range across the embeddings. We have experimented with replacing the mean pooling layer with a distance-weighted averaging function and a learned attention-based pooling layer; both approaches did increase the distribution of cosine similarities, but both also resulted in inferior performance across downstream tasks. Nonetheless, when normalized the comparisons are robust and locally specific at a resolution of one residue: less than 3% of neighboring environments would be considered significantly similar at *p* = 1 × 10^−4^.

### Supplementary Note 2

There are many hyperparameters involved in the pre-training phase, and we were not able to exhaustively search all possibilities due to computational constraints. Some parameters, including embedding dimension, environment size, and number and size of GVP layers, were therefore selected based on prior knowledge. We chose 512 for our embedding dimension because it is a middle ground between the lower-dimensional embeddings typically used in graph-based models and the higher-dimensional embeddings typically used in sequence-based models. We chose an environment size of 10 Å based on previous work with FEATURE and 3D CNNs, which has found that atomic details beyond 8–10 angstroms do not provide enough additional information to justify the additional memory and computational cost ^26, 66, 67^. Finally, the GVP architecture was adopted directly from Jing et al. (2021). ^41^ There are also alternative graph featurization schemes which could be used, including those that incorporate physico-chemical features directly. We have experimented with the addition of a feature denoting whether an edge represents a covalent bond, but it didn’t have much effect on the outcome of the model. We expect that such features are inferred by the model during training based on the raw atom types and positions, as is commonly observed of deep learning algorithms in the high-data regime. Nonetheless, it is possible that the inclusion of certain additional properties as features during the pre-training phase could improve the quality of the embeddings.

### Supplementary Note 3

#### Protein Interface Prediction (PIP)

The PIP dataset contains protein-protein interactions mined from the PDB and split by 30% sequence identity. The task is set up as a binary classification of whether or not a pair of residues, one from each interacting chain, are in contact in the bound interface. For each pair, we embed the environments around each residue separately and concatenate the embeddings. We then train a feed-forward neural network on the combined embeddings to predict whether the residues are in contact. We use one hidden layer with dimension 2048, followed by ReLU activation and dropout with 50% probability.

#### Mutation Stability Prediction (MSP)

The MSP dataset consists of pairs of wild-type and mutant protein complexes, split by 30% sequence identity. The task is set up as a binary classification of whether or not the introduction of the mutation increases or decreases the stability of the complex. Like PIP, we embed the environments around each residue in the pair, concatenate, and train a feed-forward network to predict the binary outcome.

### Supplementary Note 4

We retrieved all PDBs and labeled surface points from the MaSIF repository (https://github.com/LPDI-EPFL/masif) for the train and test splits evaluated in Gainza et al. ^47^ as well as the more difficult test set of transient interactions. For all datasets, we computed the nearest residue to the coordinates of each surface point and computed a COLLAPSE embedding of the environment centered around that residue. The prediction model consisted of the COLLAPSE encoder and a feed-forward prediction head which comprised two hidden layers of dimension 1024 with ReLU activation and dropout with probability 0.1. We report overall AUROC across all residues in each test set and compare to the corresponding metrics from Gainza et al. While our model performs slightly worse than MaSIF-site, we note that our method is designed to represent sites centered around residues rather than surfaces centered around sampled surface points. This means that the number of examples seen by our model in training is reduced by multiple orders of magnitude relative to MaSIF (158 ± 65 residues vs. 3906 ± 1428 surface points), which may account for the difference in performance.

### Supplementary Note 5

We compared the performance of COLLAPSE to analogous methods built using the large language model embeddings from ESM-1b ^50^. We computed embeddings for each site in our Prosite dataset and implemented the same training and evaluation procedures for site-specific predictions. We find that when trained in cross-validation, ESM-1b embeddings can almost perfectly classify the TP examples. This is likely due to the fact that these examples are selectd based on sequence motifs, which are easily picked up by the language model. When evaluated on the FN and FP proteins, results are more mixed; some sites perform better than COLLAPSE and some perform worse. Most notably, for IG_MHC, the ESM-1b embeddings are unable to identify any of the FN examples. We also evaluated the ability of ESM-1b embeddings to conduct local functional site searches and functional site annotation, where they generally underperform COLLAPSE embeddings in both cases. In summary, language model embeddings have strengths that largely complement structural embeddings such as COLLAPSE, and some tasks may be better suited for one than the other.

### Supplementary Note 6

As an additional baseline, we evaluated the ability of MMSeqs2 ^68^, a widely used method for sequence-based searches, to reclassify Prosite FN and FP proteins. For each sequence annotated as a FN or FP, we conducted a single search against the Swissprot database. We then extracted the top hit from this search and checked whether this protein was a known example of that particular functional site. If so, we consider that protein successfully re-classified. The results shown in Table S4 clearly demonstrate the value of training site-specific prediction models and of the use of local structural embeddings, as very few of the FN proteins are successfully identified as members of the functional site. However, our simple MMSeqs2 baseline is very effective at ruling out FPs.

### Supplementary Note 7

The procedure for computing FEATURE vectors for the MSP dataset was as follows:

1. Create two PDB files for every example in the dataset, one for wild-type and one for mutated.
2. Compute structural features of each PDB using DSSP ^69^.
3. Create two .ptf files for every example in the dataset, one for wild-type and one for mutated, using the coordinates of the functional atom of the central residue.
4. Compute FEATURE vectors using *featurize* from the FEATURE package (https://simtk.org/projects/feature; see user manual for details).

The training procedure is identical to that used for the COLLAPSE benchmark. While FEATURE performs quite well relative to the ATOM3D benchmarks on this difficult task, COLLAPSE still outperforms it when fine-tuned. We note that the inability to fine-tune FEATURE on task-specific datasets is a major limitation which is addressed by our self-supervised learning framework. Unfortunately, we were not able to fully train a FEATURE benchmark for PIP, since the computation of FEATURE vectors is not very efficient for very large datasets. Given the relative difficulty of using FEATURE relative to COLLAPSE, its inability to be fine-tuned, and its inferior performance across many benchmarks, we are confident that COLLAPSE provides an improved general-purpose representation of protein sites.

### Supplementary Note 8

We obtained all PDBs and site data for each dataset from the following URL https://www.ccs.neu.edu/home/radivojac/data/xin_cpps_supplementary.zip. We then embedded all residues in each dataset and split the embeddings randomly into 10 folds, as in Xin et al. ^48^ The model and training protocol was exactly the same as for the Prosite models (Section 4.4): we train an SVM with radial basis function kernel and evaluate total AUROC over all held-out folds. We compare against the three methods presented in Xin et al., without retraining. Note that the exact splits for each fold are different between our model and the baseline models.

### Supplementary Note 9

We obtained the train, validation, and test splits for both remote homology and enzyme classification tasks from Hermosilla et al., ^70^ downloaded the corresponding PDB chains and domains, and embedded all residues in each. All ligands and heteroatoms were excluded. We then trained a bidirectional two-layer gated recurrent unit (GRU) model to aggregate over the full sequence of embeddings for each protein. The GRU output was computed as the concatenation of the final hidden state of the forward and backward networks in the last layer, which was fed into a two-layer feed-forward network with ReLU activation and dropout with probability 0.5 to predict the final classification. Basic hyperparameter tuning was performed, and the best model was selected based on the validation accuracy. The final fold prediction model used a learning rate of 1 × 10^−3^, batch size of 64, hidden unit size of 256, and an AdamW optimizer with weight decay of 0.5. The enzyme class prediction model used a learning rate of 1 × 10^−4^, batch size of 32, hidden unit size of 1024, and an AdamW optimizer with weight decay of 0.5. We compare to all baseline methods presented in Hermosilla et al., including the hidden Markov model (HMM) based method HHSuite as well as sequence- and structure-based machine learning models, many of which were fine-tuned specifically for each task. Our model, on the other hand, only aggregates over fixed COLLAPSE embeddings. Notably, HHSuite is one of the top performers for all tasks except the most difficult homology split, where we achieve much higher accuracy. The performance of the other methods varies across tasks, and even without fine-tuning our COLLAPSE-RNN model consistently ranks in the top five methods for all tasks and test sets. We expect that this performance could be improved by fine-tuning or by improving the aggregation method from the simple RNNs used here.

## Supplementary Figures

**Figure S1.**
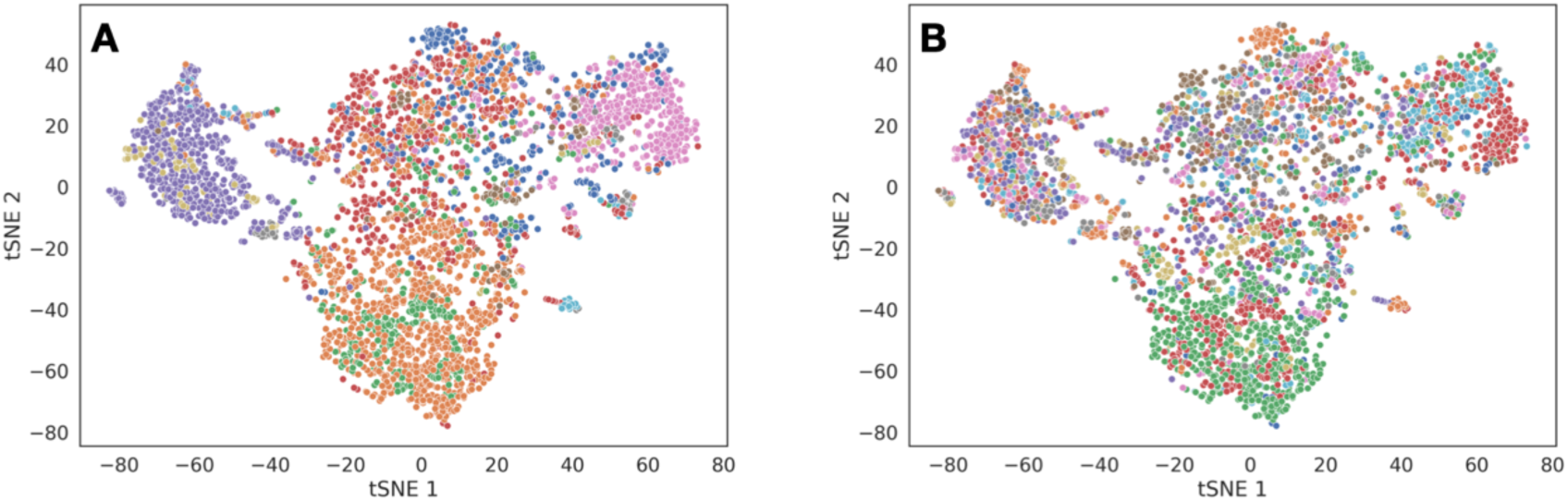
PCA of embeddings of single-domain proteins at lower levels of the CATH hierarchy: **(a)** architecture and **(b)** topology.

**Figure S2.**
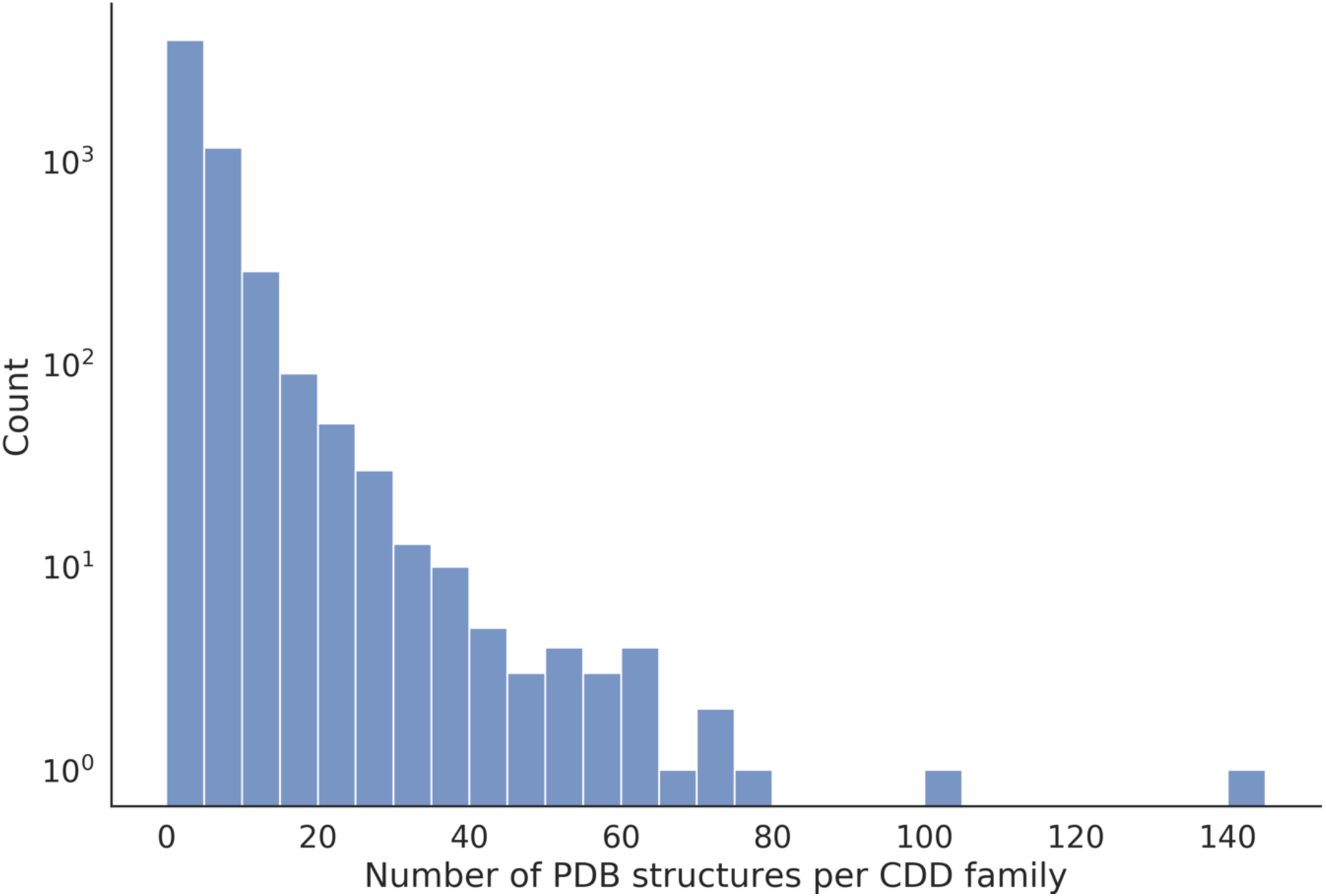
Histogram of number of PDB structures per CDD family in training dataset.

**Figure S3.**
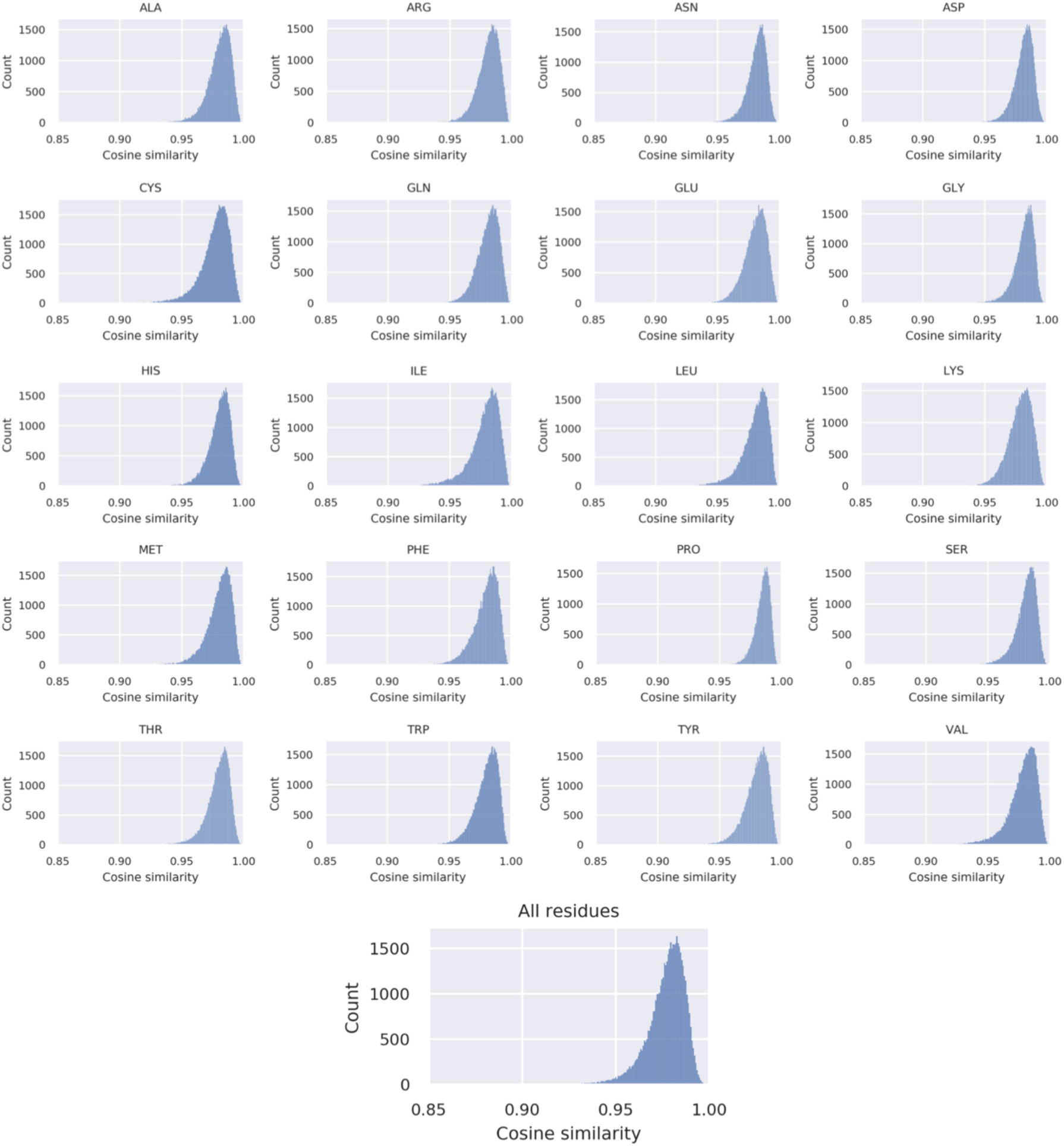
Empirical cosine similarity distributions computed for each amino acid and the combined dataset.

**Figure S4.**
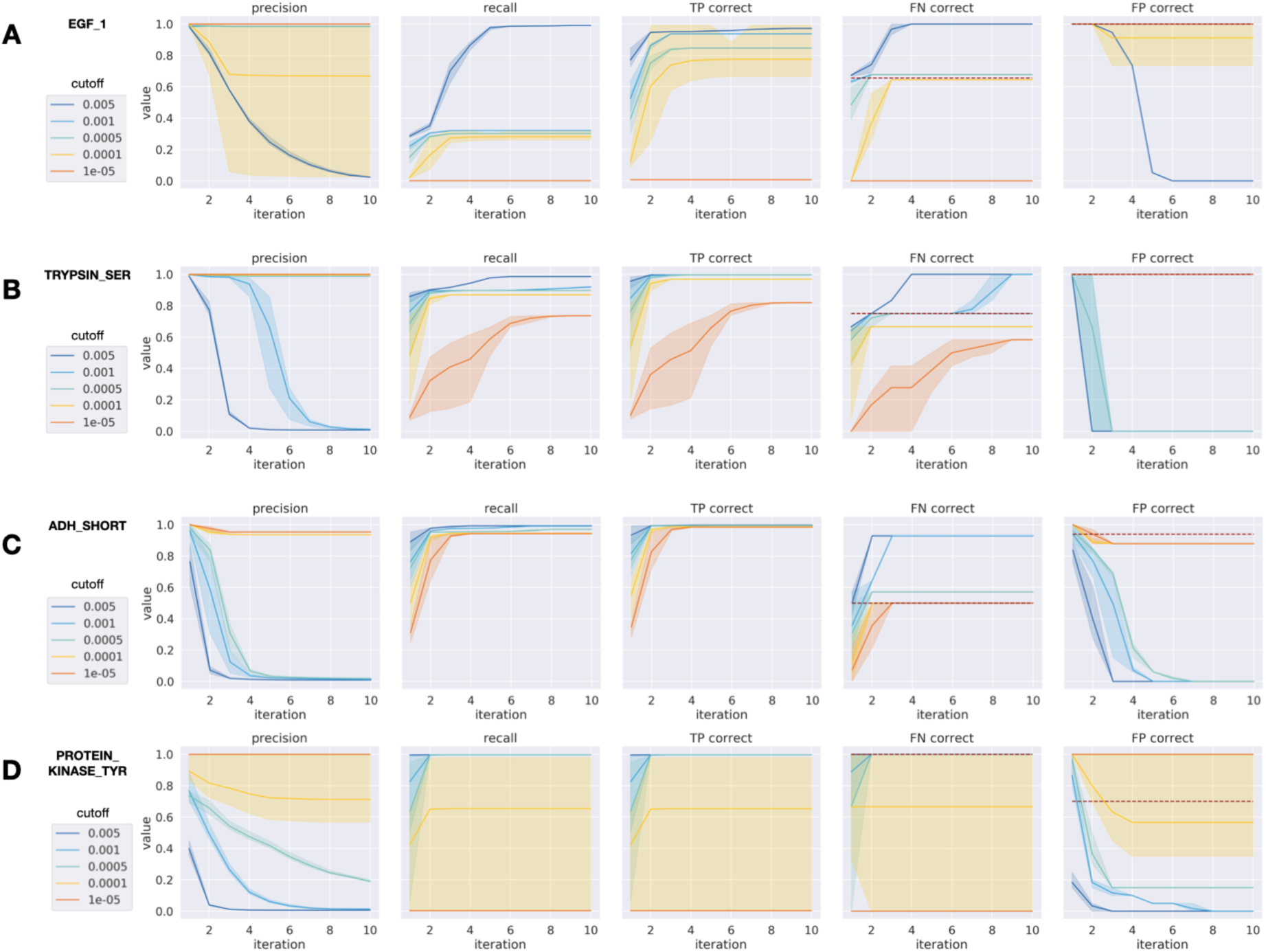
Functional site search performance per iteration for remaining Prosite families with FP and FN annotations not shown in Figure 5: **(a)** EGF_1, **(b)** TRYPSIN_SER, **(c)** ADH_SHORT, and **(d)** PROTEIN_KINASE_TYR.

**Figure S5.**
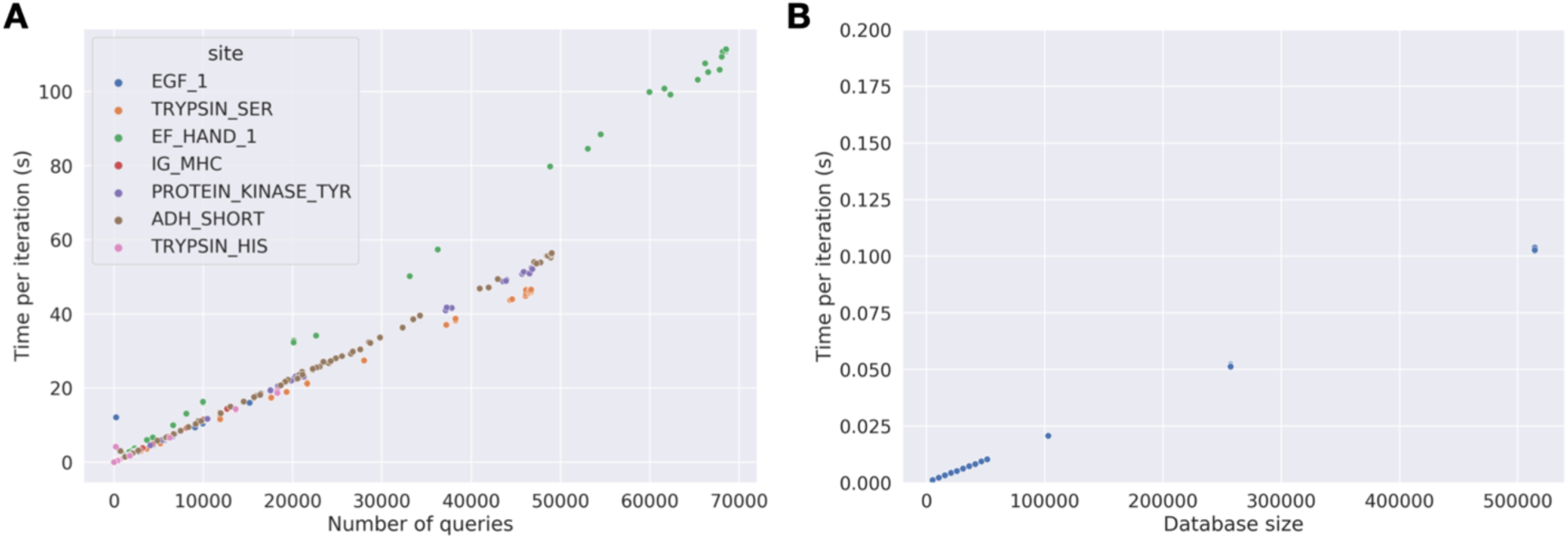
Runtime analysis for functional site search tool. Time per iteration as a function of **(a)** number of queries at the start of the iteration, colored by functional site, and **(b)** the size of the database searched against.

**Figure S6.**
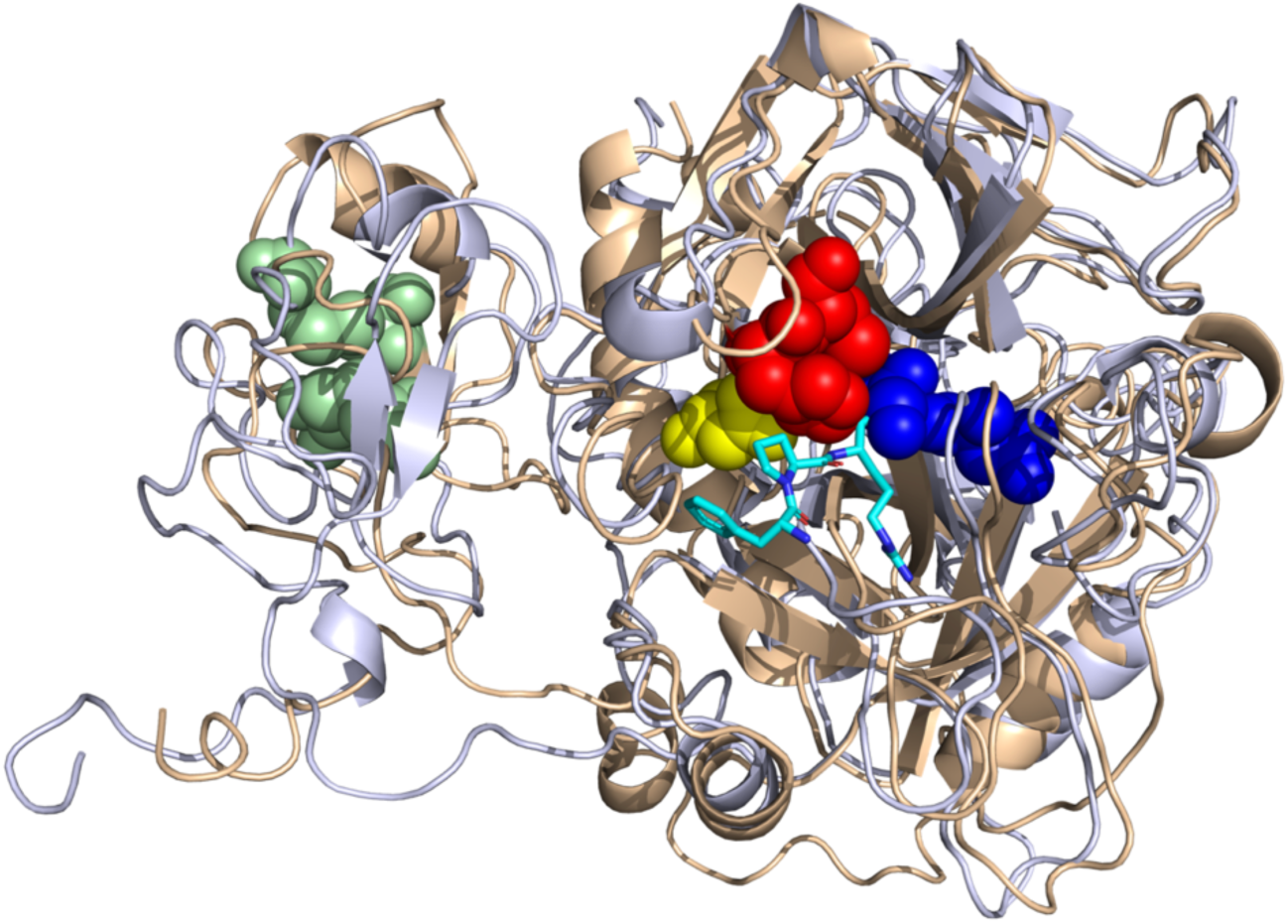
Annotated structure of meizothrombin structure predicted by AlphaFold (gold; Uniprot ID P00735) superimposed on crystal structure (light blue; PDB ID 1A0H). Colors correspond to the predicted functional site, using the same colors as Fig. 5a.

**Figure S7.**
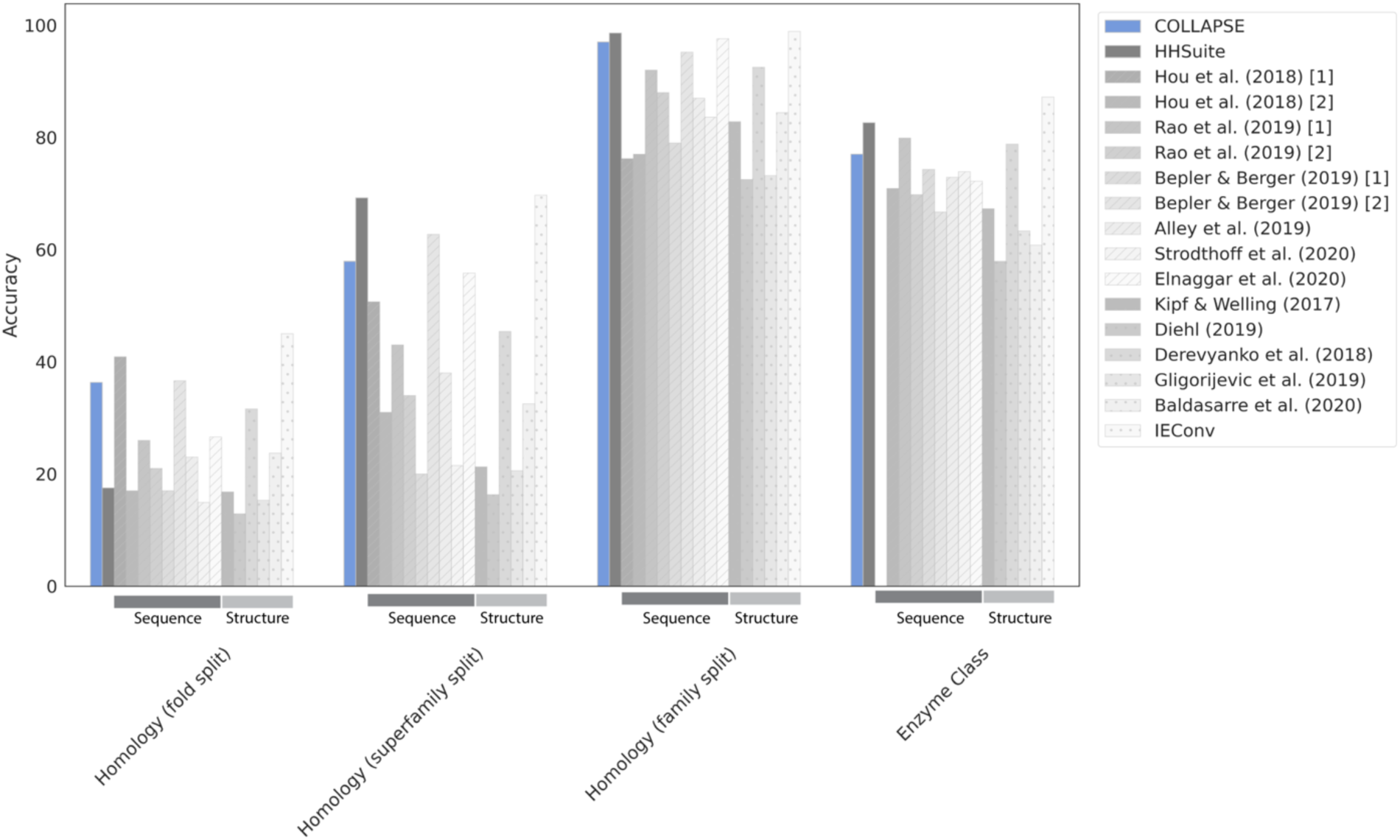
Performance of COLLAPSE on protein-level prediction tasks using a GRU aggregator. The tasks are fold prediction (homology) and enzyme class prediction using datasets and baseline methods from Hermosilla et al. (2021). ^70^. We report results for three held-out test sets for the homology task, defined by the stringency of the structural overlap with the training set, with “fold split” being the most difficult and “family split” being the easiest. Our COLLAPSE-RNN method is shown in blue, and the HMM-based HHsuite is shown in dark gray. For the ML-based baselines, we note which operate primarily on sequences and which operate primarily on structure using diagonal lines and dots, respectively.

**Figure S8.**
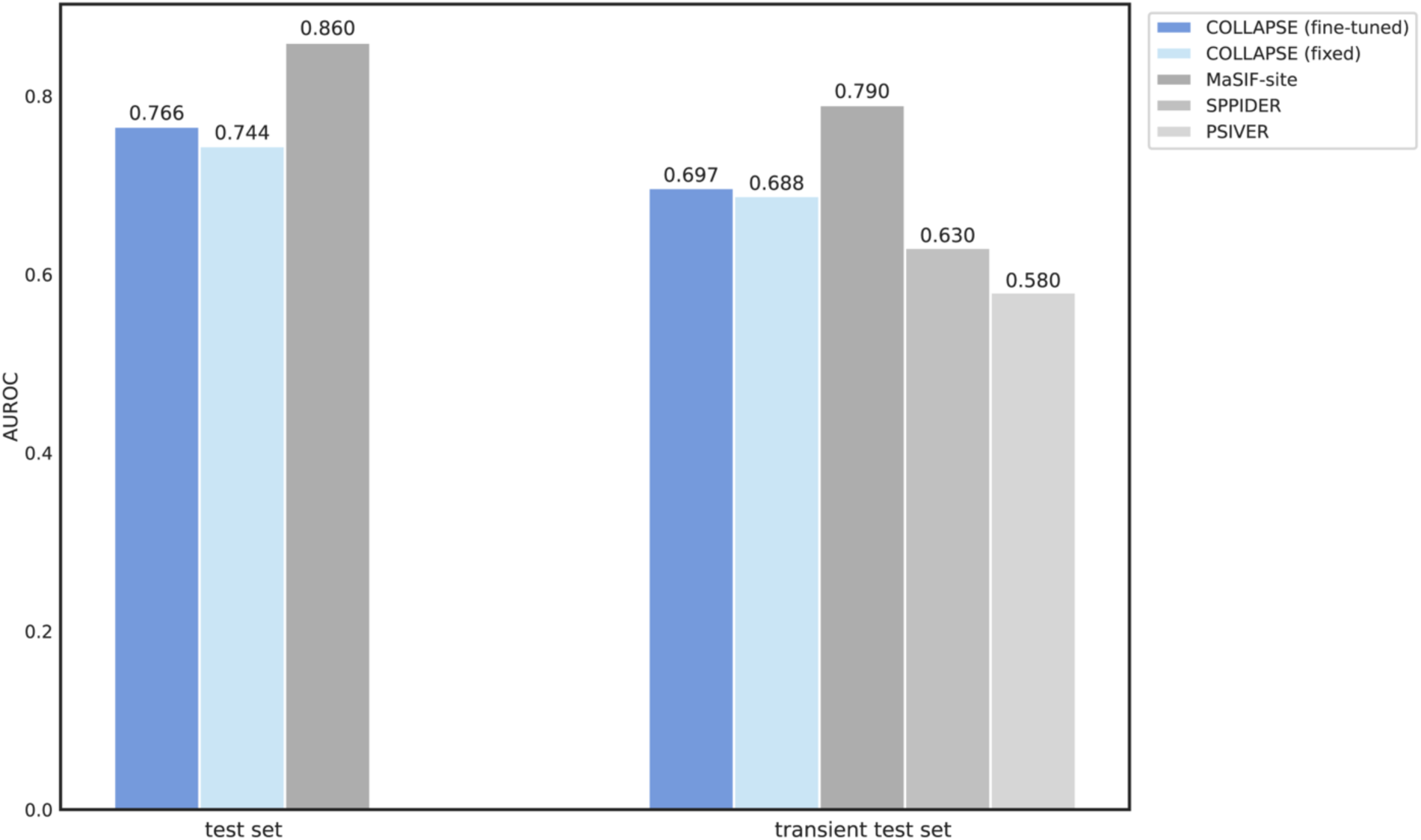
Performance of COLLAPSE on protein-protein interaction binding site identification. We compare to MaSIF-site as well as two baseline methods, as reported in Gainza et al. (2020). ^47^ The metric is AUROC over all residues in all proteins, and we evaluate on both the full test set (left) and the more difficult subset of transiently interacting proteins (right). See Supplementary Note 4 for details.

**Figure S9.**
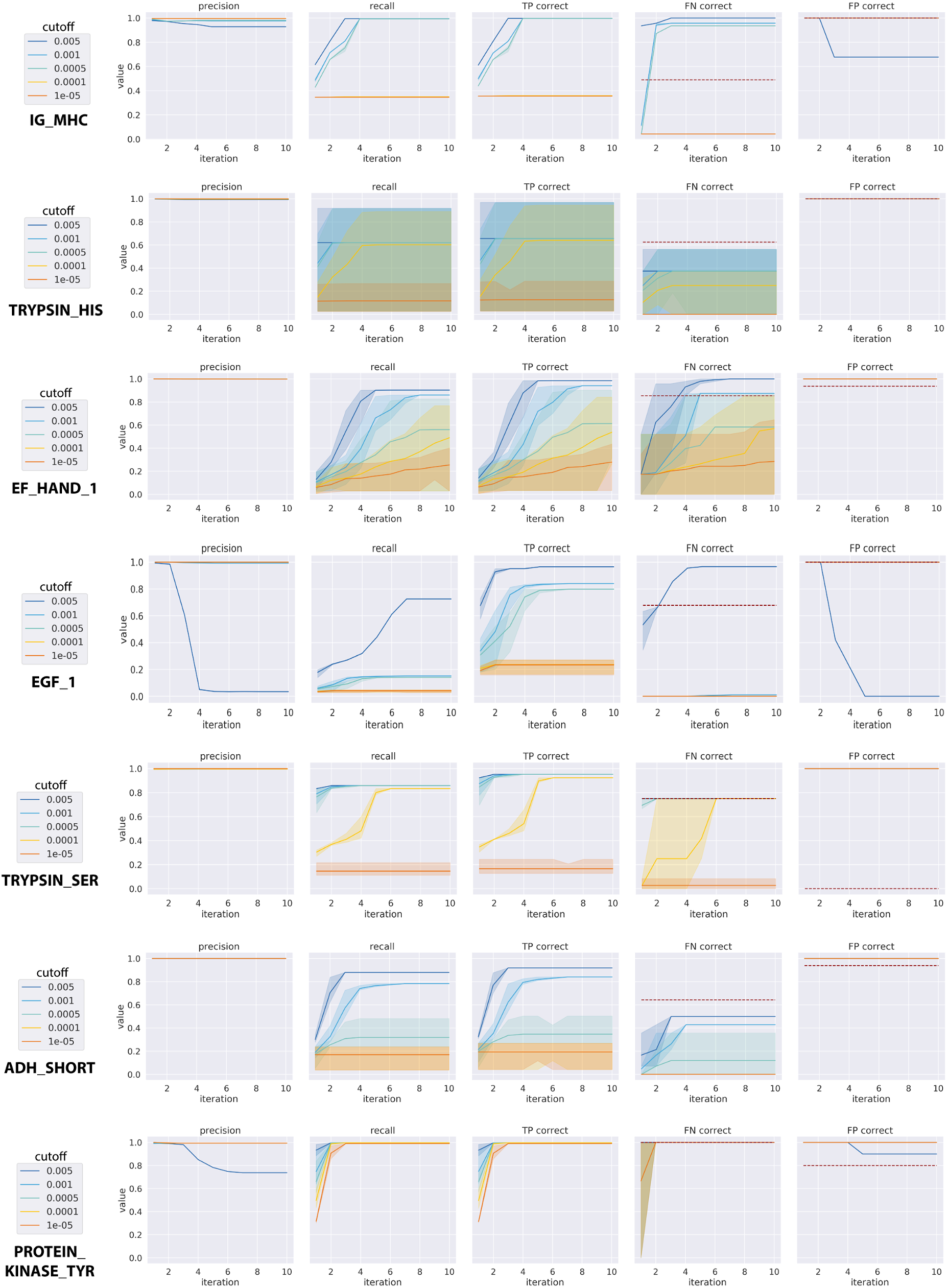
Performance of ESM-1b language model embeddings in functional site search (see Supplementary Note 5 for details). Sequence embeddings are not able to match the recall of COLLAPSE over most sites, even after many iterations, although precision is often high.

**Figure S10.**
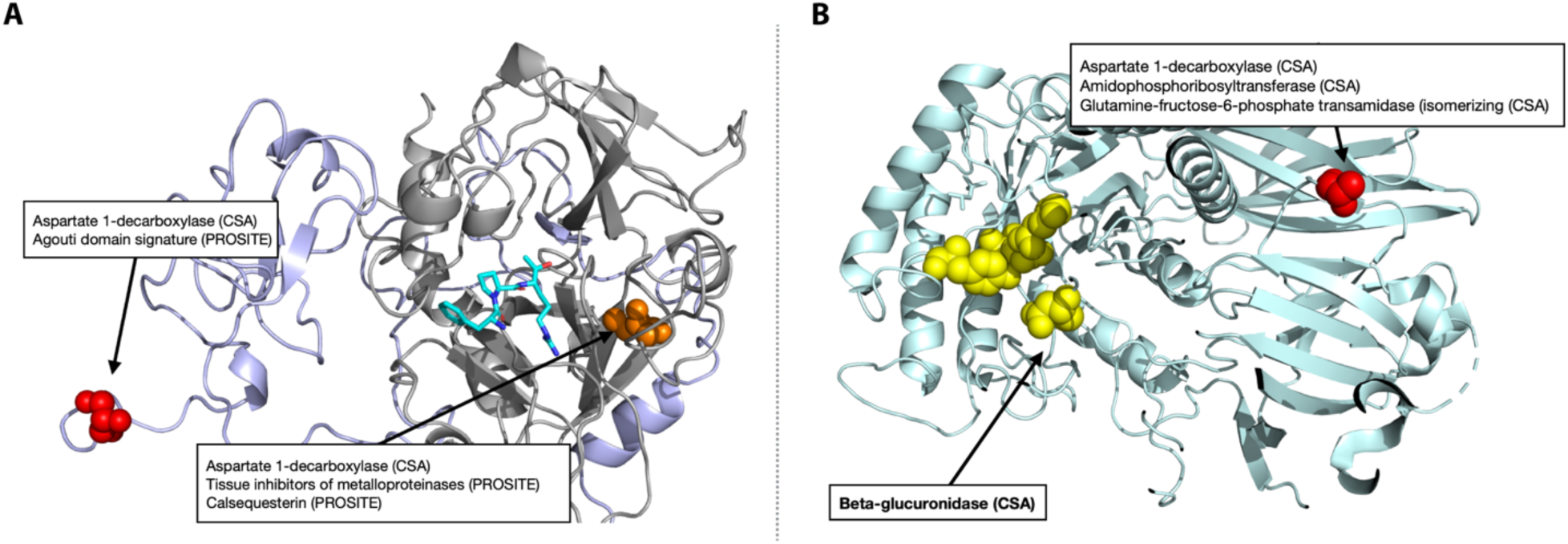
Performance of ESM-1b in functional site annotation (see Supplementary Note 5 for details) for **(a)** Meizothrombin (PDB ID: 1A0H) at *p* = 5 × 10^−5^ and **(b)** beta-glucuronidase (PDB ID: 3HN3) at *p* = 1 × 10^−4^. In both cases, ESM-1b sequence embeddings result in lower sensitivity and more false positives than COLLAPSE embeddings. For meizothrombin, there are no correct predictions at a p-value of 5 × 10^−5^. When the threshold is lowered to 1 × 10^−4^, the serine and histidine active sites and the kringle domain are recognized, but at the cost of 10 false positive annotations. The catalytic aspartic acid is not identified. For beta-glucuronidase, one of the three active sites is correctly identified at *p* = 1 × 10^−4^, but there are also three false positive annotations at a residue far from the active site.

## Supplementary Tables

**Table S1.**
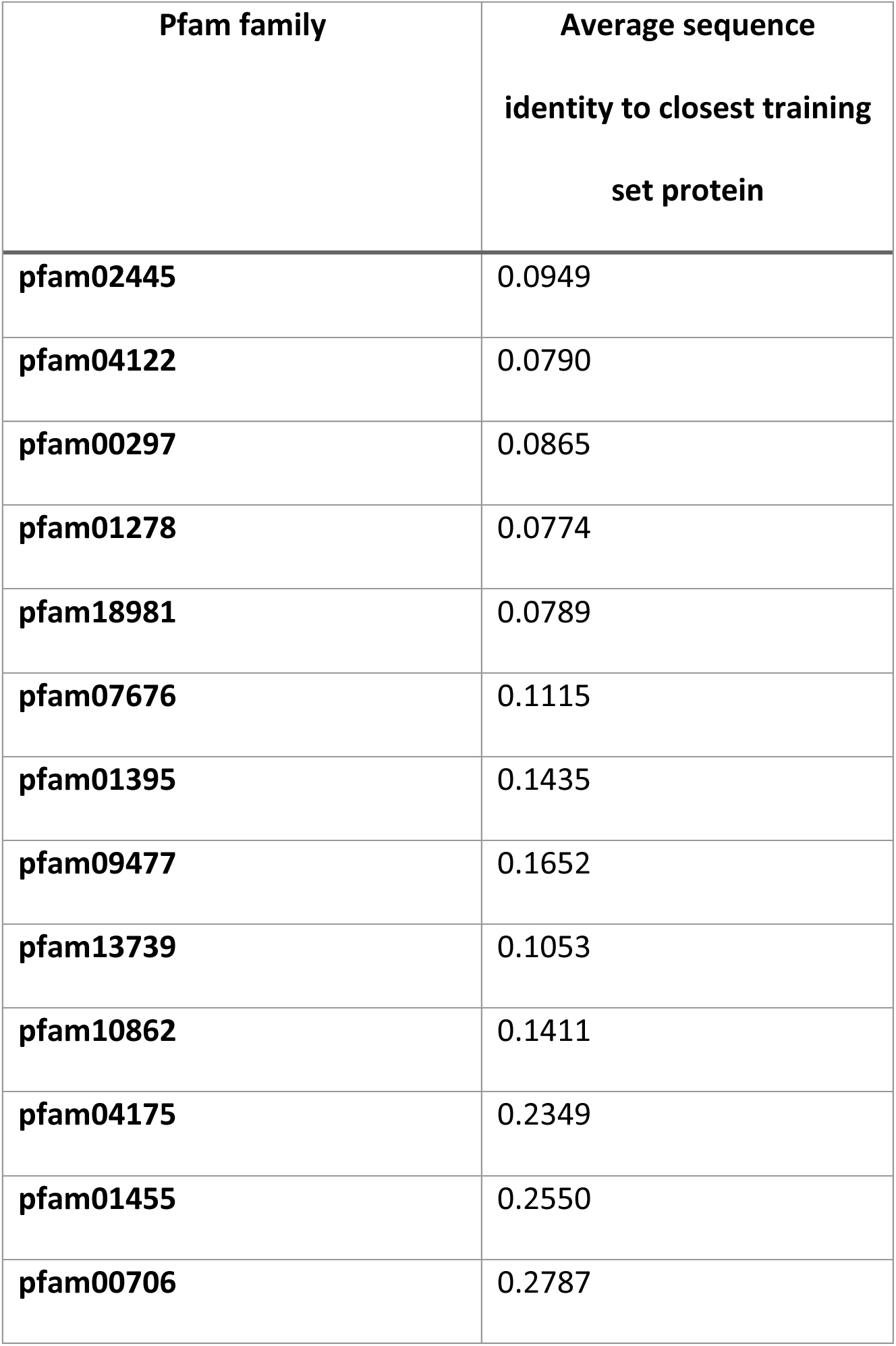
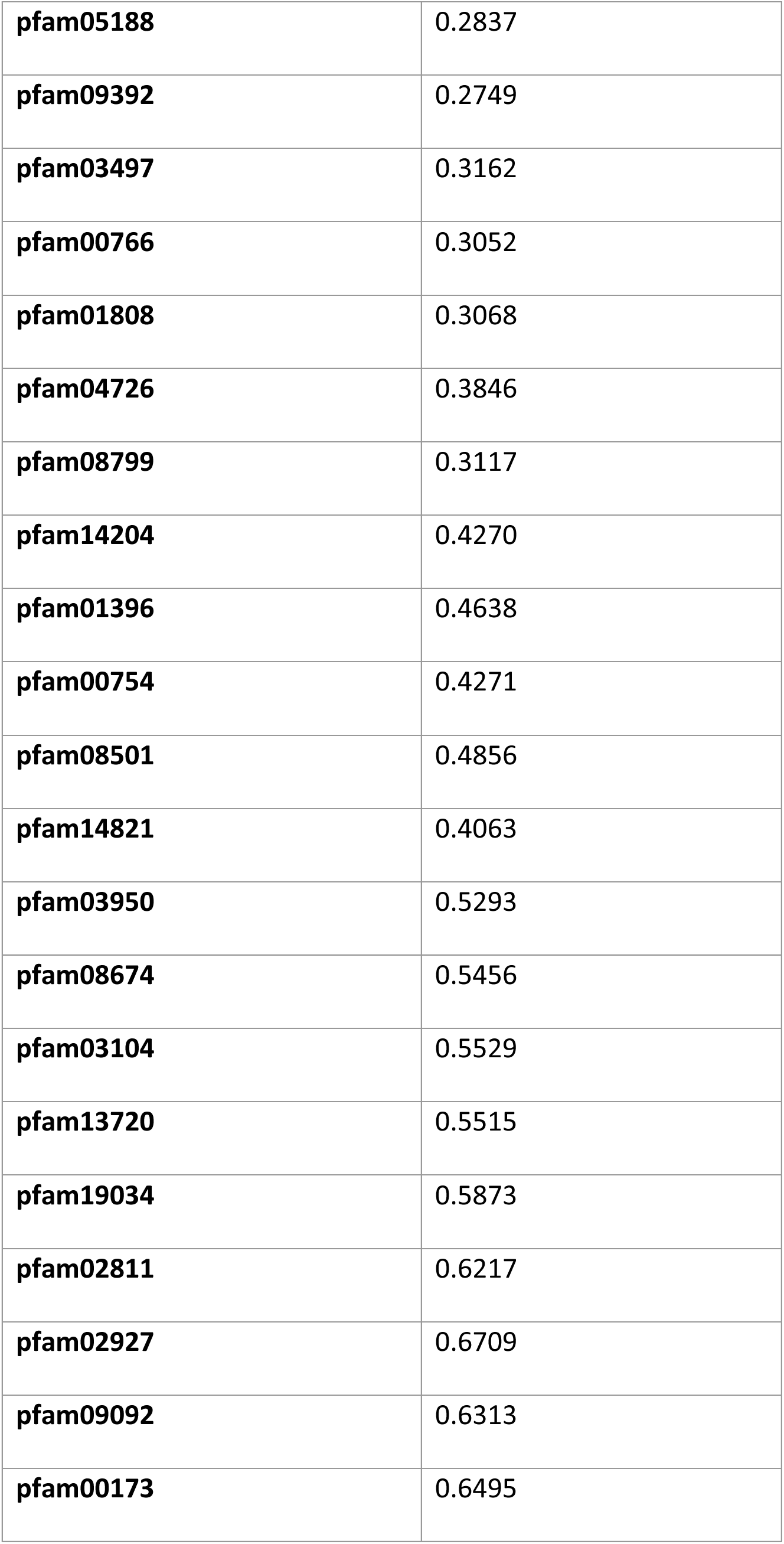
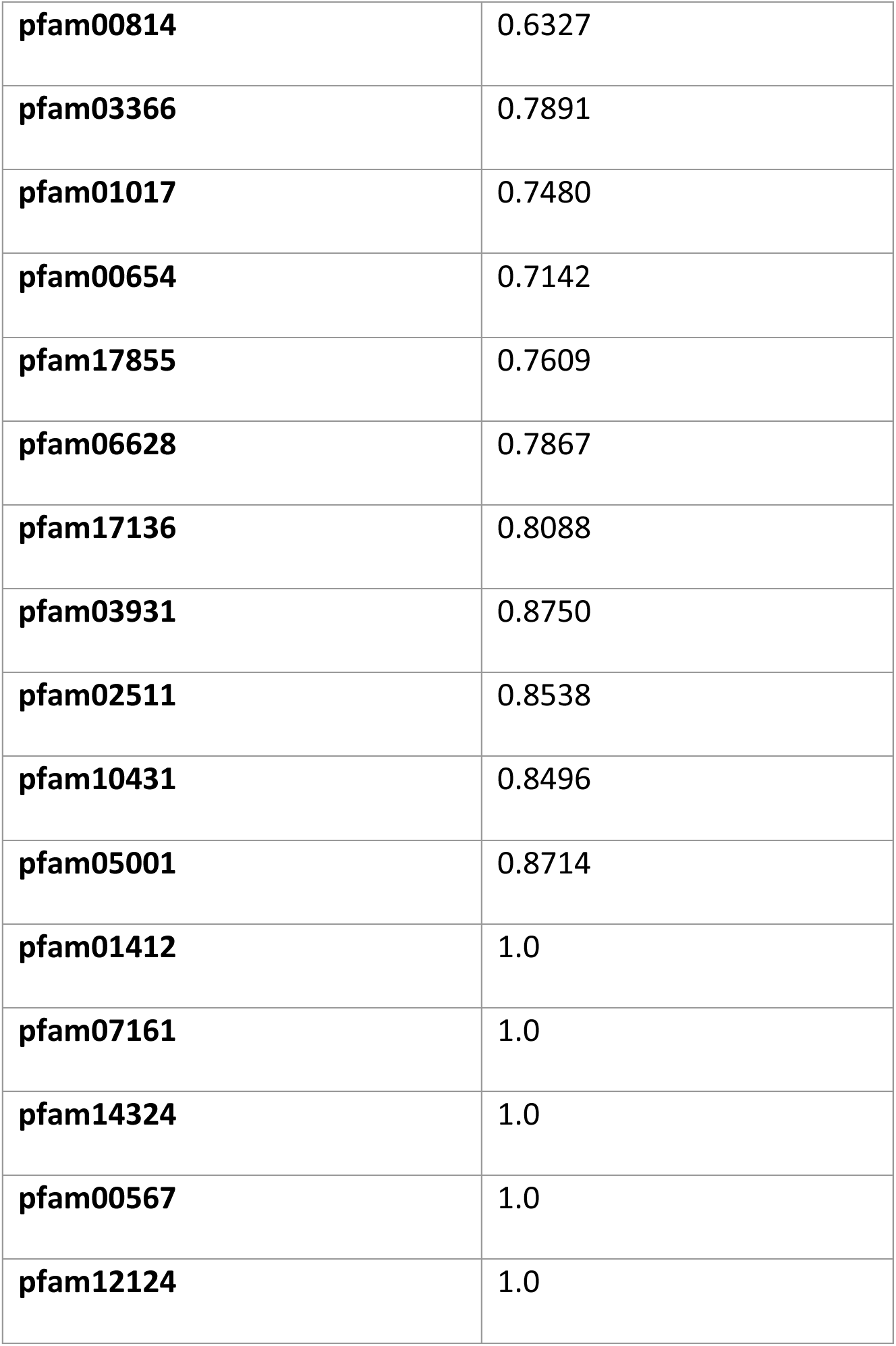
Pfam families selected for held-out validation set and corresponding sequence identity to nearest protein in CDD training set.

| **Pfam family** | **Average sequence**<br>**identity to closest training**<br>**set protein** |
| --- | --- |
| **pfam02445** | 0.0949 |
| **pfam04122** | 0.0790 |
| **pfam00297** | 0.0865 |
| **pfam01278** | 0.0774 |
| **pfam18981** | 0.0789 |
| **pfam07676** | 0.1115 |
| **pfam01395** | 0.1435 |
| **pfam09477** | 0.1652 |
| **pfam13739** | 0.1053 |
| **pfam10862** | 0.1411 |
| **pfam04175** | 0.2349 |
| **pfam01455** | 0.2550 |
| **pfam00706** | 0.2787 |
| **pfam05188** | 0.2837 |
| **pfam09392** | 0.2749 |
| **pfam03497** | 0.3162 |
| **pfam00766** | 0.3052 |
| **pfam01808** | 0.3068 |
| **pfam04726** | 0.3846 |
| **pfam08799** | 0.3117 |
| **pfam14204** | 0.4270 |
| **pfam01396** | 0.4638 |
| **pfam00754** | 0.4271 |
| **pfam08501** | 0.4856 |
| **pfam14821** | 0.4063 |
| **pfam03950** | 0.5293 |
| **pfam08674** | 0.5456 |
| **pfam03104** | 0.5529 |
| **pfam13720** | 0.5515 |
| **pfam19034** | 0.5873 |
| **pfam02811** | 0.6217 |
| **pfam02927** | 0.6709 |
| **pfam09092** | 0.6313 |
| **pfam00173** | 0.6495 |
| **pfam00814** | 0.6327 |
| **pfam03366** | 0.7891 |
| **pfam01017** | 0.7480 |
| **pfam00654** | 0.7142 |
| **pfam17855** | 0.7609 |
| **pfam06628** | 0.7867 |
| **pfam17136** | 0.8088 |
| **pfam03931** | 0.8750 |
| **pfam02511** | 0.8538 |
| **pfam10431** | 0.8496 |
| **pfam05001** | 0.8714 |
| **pfam01412** | 1.0 |
| **pfam07161** | 1.0 |
| **pfam14324** | 1.0 |
| **pfam00567** | 1.0 |
| **pfam12124** | 1.0 |

**Table S2.**
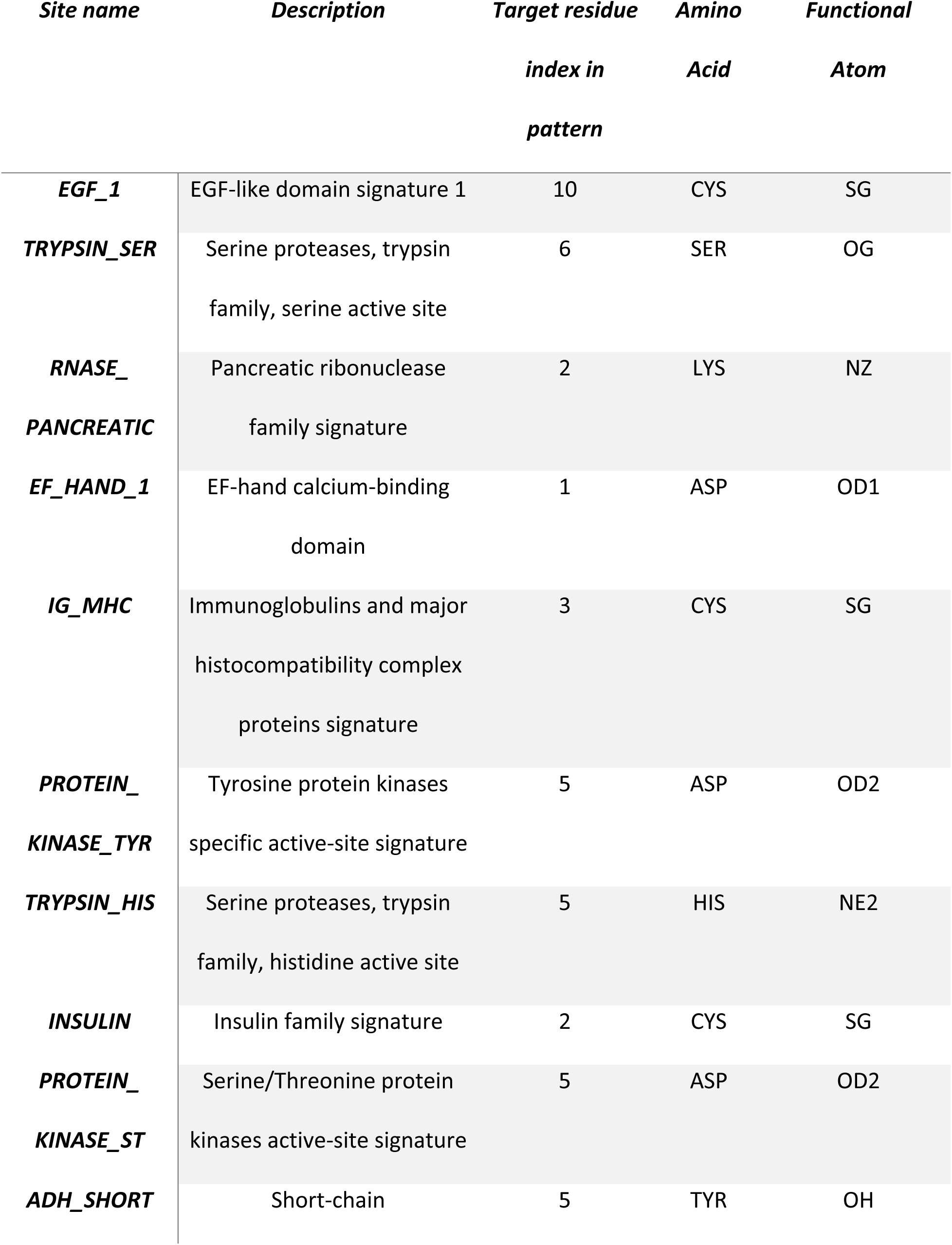
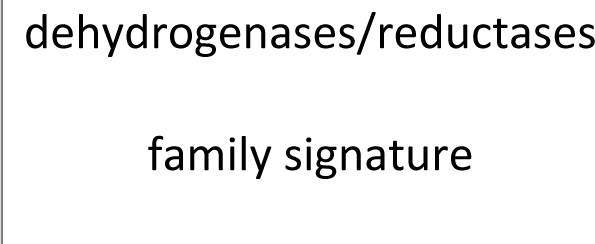
Description of 10 Prosite functional sites and definition of functional centers for each.

| *Site name* | *Description* | *Target residue*<br>*index in*<br>*pattern* | *Amino*<br>*Acid* | *Functional*<br>*Atom* |
| --- | --- | --- | --- | --- |
| **EGF_1** | EGF-like domain signature 1 | 10 | CYS | SG |
| **TRYPSIN_SER** | Serine proteases, trypsin<br>family, serine active site | 6 | SER | OG |
| **RNASE_**<br>**PANCREATIC** | Pancreatic ribonuclease<br>family signature | 2 | LYS | NZ |
| **EF_HAND_1** | EF-hand calcium-binding<br>domain | 1 | ASP | OD1 |
| **IG_MHC** | Immunoglobulins and major<br>histocompatibility complex<br>proteins signature | 3 | CYS | SG |
| **PROTEIN_**<br>**KINASE_TYR** | Tyrosine protein kinases<br>specific active-site signature | 5 | ASP | OD2 |
| **TRYPSIN_HIS** | Serine proteases, trypsin<br>family, histidine active site | 5 | HIS | NE2 |
| **INSULIN** | Insulin family signature | 2 | CYS | SG |
| **PROTEIN_**<br>**KINASE_ST** | Serine/Threonine protein<br>kinases active-site signature | 5 | ASP | OD2 |
| **ADH_SHORT** | Short-chain | 5 | TYR | OH |

|  |
| --- |
| dehydrogenases/reductases |
| family signature |

**Table S3.** Dataset statistics for Prosite binary classification task.

| *Dataset* | *Positive Examples* | *Negative Examples* |
| --- | --- | --- |
| *EGF_1* | 144 | 50,023 |
| *TRYPSIN_SER* | 315 | 48,032 |
| *RNASE_PANCREATIC* | 393 | 49,996 |
| *EF_HAND_1* | 1,971 | 48,141 |
| *IG_MHC* | 2,188 | 49,073 |
| *PROTEIN_KINASE_TYR* | 301 | 50,004 |
| *TRYPSIN_HIS* | 445 | 47,476 |
| *INSULIN* | 438 | 49,074 |
| *PROTEIN_KINASE_ST* | 1,199 | 49,999 |
| *ADH_SHORT* | 383 | 50,128 |

**Table S4.**
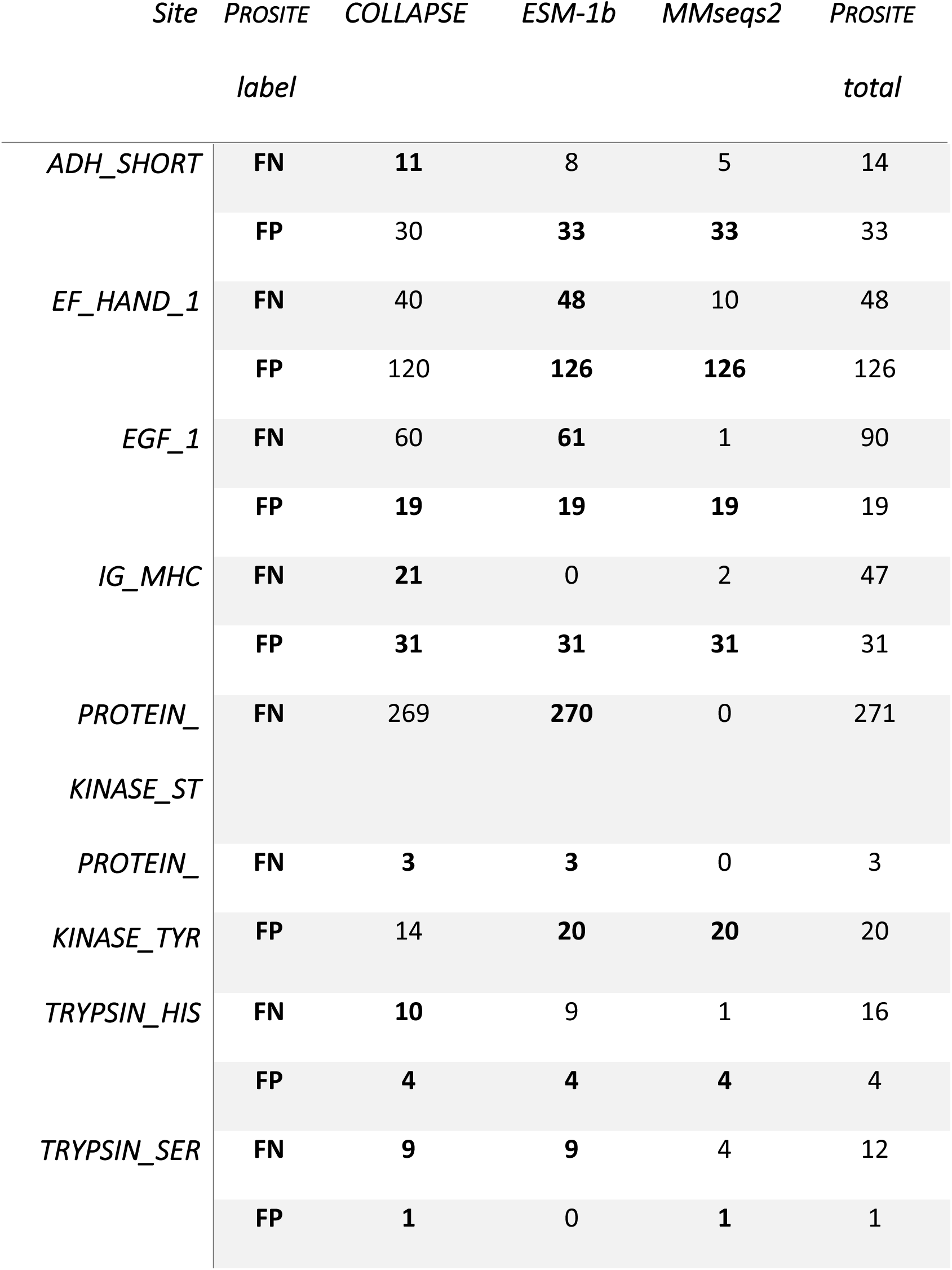
Performance on Prosite FN/FP proteins compared to ESM-1b language model embeddings ^50^ and MMseqs2. ^68^ See Supplementary Notes 5–6 for details.

**Table S5.** Performance of COLLAPSE on functional site identification benchmarks from Xin et al. (2011). ^48^ Metric is AUROC, and highest-performing method is in bold. Performance of baseline methods is taken directly from the paper, so the exact splits used in each fold are not identical. See Supplementary Note 8 for details.

|  | *Zn-binding* | *DNA-binding* | *Catalytic* | *Phosphorylation* |
| --- | --- | --- | --- | --- |
| *COLLAPSE* | **0.884** | 0.812 | **0.889** | 0.647 |
| *FEATURE* | 0.767 | **0.824** | 0.754 | 0.620 |
| *GBT* | 0.713 | 0.713 | 0.814 | 0.559 |
| *Graphlet kernel* | 0.758 | 0.619 | 0.732 | 0.688 |
| *Structure kernel* | 0.808 | 0.800 | 0.839 | **0.711** |

**Table S6.** ATOM3D benchmarking results compared to FEATURE for MSP dataset. The metric is area under the receiver operator characteristic curve (AUROC), and we report mean and standard deviation across three training runs. Numbers in bold indicate best performance on each task (within one standard deviation). See Supplementary Note 7 for details.

| *Task (metric)* | *COLLAPSE (fixed)* | *COLLAPSE (fine-tuned)* | *FEATURE (fixed)* |
| --- | --- | --- | --- |
| *MSP (AUROC)* | 0.616 ± 0.006 | **0.668 ± 0.018** | 0.617 ± 0.012 |

